# The Limits of Molecular Signatures for Pancreatic Ductal Adenocarcinoma Subtyping

**DOI:** 10.1101/2022.05.16.491983

**Authors:** Manuela Lautizi, Jan Baumbach, Wilko Weichert, Katja Steiger, Markus List, Nicole Pfarr, Tim Kacprowski

## Abstract

Molecular signatures have been suggested as biomarkers to classify pancreatic ductal adenocarcinoma (PDAC) into two, three or four subtypes. Since the robustness of existing signatures is controversial, we performed a systematic evaluation of three established signatures for PDAC stratification across eight publicly available datasets. Clustering revealed inconsistency of subtypes across independent datasets and in some cases a different number of PDAC subgroups than in the original study, casting doubt on the actual number of existing subtypes. Next, we built nine classification models to investigate the ability of the signatures for tumor subtype prediction. The overall classification performance ranged from ∼35% to ∼90% accuracy, suggesting instability of the signatures. Notably, permuted subtypes and random gene sets achieved very similar performance. Cellular decomposition and functional pathway enrichment analysis revealed strong tissue-specificity of the predicted classes. Our study highlights severe limitations and inconsistencies that can be attributed to technical biases in sample preparation and tumor purity, suggesting that PDAC molecular signatures do not generalize across datasets. How stromal heterogeneity and immune compartment interplay in the diverging development of PDAC is still unclear. Therefore, a more mechanistic or at least multi-omic approach seems necessary to extract more robust and clinically exploitable insights.

## INTRODUCTION

### Background and context

Pancreatic Ductal Adenocarcinoma (PDAC) is the most frequent and aggressive malignant neoplasm of the pancreas. Its incidence has been increasing in the past decades and it is expected to rise further, reinforcing PDAC’s position as one of the deadliest cancer types and on the way of becoming the second leading cause of cancer-related death by 2030 [1]. The absence of clear symptoms leads to a delayed diagnosis of the disease, where the tumor stage is often advanced and traditional treatments do not necessarily prolong the survival time. Around half of the patients show spread of distance metastases already at an early stage or when the tumor has still a small diameter (< 2 cm), which renders surgical resection and radiotherapy no longer applicable [2,3]. Despite progress in anti-cancer drug research, chemotherapy yields only minor survival advantages while strongly affecting the quality of life due to its high toxicity [4]. More recently, immunotherapy has shown encouraging treatment results. However, the PDAC tumor microenvironment is widely heterogeneous and characterized by the presence of immunosuppressive cells, which, in a dense stroma environment, makes the malignancy resistant to immunotherapy as well as chemotherapy [5–7].

Precision medicine, i.e. the stratification of patients into clinically actionable cancer subtypes based on molecular characteristics of a tumor, is expected to significantly impact the outcome response [8]. Different cancer subtypes present different mechanisms involved in carcinogenesis and are linked to different phenotypes [9]. At the same time, patients within the same subgroup share common molecular patterns and genomic alterations, as well as similar clinical outcomes. Widely available transcriptomics data is particularly attractive for defining molecular subtypes as was initially demonstrated successfully in the context of breast cancer, where gene expression profiling drives subgroup classification [10,11]. In the same way, researchers proposed the identification of molecular subtypes in PDAC.

### Molecular subtypes of pancreatic cancer

Many studies built up the current knowledge of PDAC molecular subtypes, proposing a tumor classification which reflects the biological and prognostic differences between patients. Here, we focus on the three transcriptomic and genomic studies most frequently discussed in literature for the stratification of PDAC into different groups. These studies are the ones conducted by Collisson *et al*. [12], Moffitt *et al*. [13] and Bailey *et al*. [14], which are used as a gold standard for PDAC molecular subtyping. After a first definition of subtypes was established by Collisson *et al*. in 2011, Moffitt *et al*. and Bailey *et al*. proposed different molecular subtypes of pancreatic cancer using larger cohorts, in 2015 and 2016, respectively.

Collisson *et al*. performed an unsupervised analysis on the combination of two microarray datasets, one coming from microdissected primary tumor samples (where the epithelium was devoid of the stroma) and a second one from whole tumor samples. The authors identified a signatures of 62 genes for patients discrimination and proposed three subtypes: Classical, Quasi-Mesenchymal (QM-PDA) and Exocrine-like. To validate these subtypes, resp. cell lines were tested for therapy response *in vitro*. The use of Gemcitabine and Erlotinib in human PDAC cell lines of known subtypes had different effects on the groups. Classical subtypes appeared to benefit more from Erlotinib, opposed to QM-PDA where Gemcitabine was more beneficial. In a validation of the 62 genes via unsupervised analysis on human and mouse cell lines, the Exocrine-like subtype was not observed. Furthermore, three additional publicly available expression datasets with corresponding survival data were used for subtype partitioning based on the 62 genes. Clustering these additional datasets together with their original data reproduced the three subtypes, whereas clustering the additional datasets individually lead to different clusters, suggesting that an independent reproduction of the subtypes in different datasets is challenging.

Moffitt *et al*. performed a virtual microdissection on the integration of multiple microarray data (primary and metastatic tumors, cell lines, normal pancreas and distant site adjacent normal samples). The transcripts were divided into categories linked with each input phenotype, and transcripts associated with tumor and stroma were used to define two distinct subtyping approaches. Tumor-specific subtypes rely on 50 tumor-related genes which stratify the patients into Classical and Basal-like subtypes, while 48 stroma-related genes were used to distinguish between normal and activated stroma subtypes. Survival analysis was used to assess how the subtypes differ prognostically. The discovered subtypes were further validated by sequencing of primary PDAC, patient derived xenografts, cell lines and cancer associated fibroblasts. While clustering on patient-derived xenografts supports the Basal/Classical classification, cell line models exhibit a prevalence of the Basal-like subtype, implying cell lines are not suitable for such binary PDAC classification.

Bailey *et al*. used RNASeq data from pancreatic cancer samples of different histopathological subtypes that have more than 40% of tumor cellularity. *Via* clustering, they identified four clusters of subtypes: Squamous, Immunogenic, Pancreatic Progenitor and Aberrantly Differentiated Exocrine (ADEX) which showed significantly different prognosis. To elucidate other biologically relevant characteristics of the subtypes, Bailey *et al*. integrated the transcriptomic analysis with a genomic analysis comprising whole-genome, deep-exome sequencing and gene copy number variation. A multiclass Significance Analysis of Microarrays (SAM) analysis returned a list of 613 genes differentially expressed between the four subtypes. Bailey *et al*. employed a larger human cohort to assess the reproducibility of the classes, this time array-based and without assessing tumor cellularity. After submitting the mRNA expression profiles to the same clustering procedure, the authors report the discovery of the same tumor subclasses.

**Table 1.** Overview on the PDAC molecular subtypes.

| Bailey | Pancreatic Progenitor | Squamous | ADEX | Immunogenic |
| --- | --- | --- | --- | --- |
|  | Transcription factors PDX1, MNX1, HNF1G, FOXA1, HES1 related to early pancreatic development and associated with fatty acid oxidation, steroid hormone biosynthesis, drug metabolism and glycosylation of mucins.<br>*Prognosis:* 23.7 survival months | Overexpression of genes implicated in inflammation, hypoxia, metabolism, activated MYC pathway, TGF-β signaling, autophagy, cell proliferation. Activated $\alpha 6 \beta 1$, $\alpha 6 \beta 4$ and EGF signaling. Samples hypermethylated with consequent downregulation of endodermal cell fate genes causing loss of endodermal identity. TP53 mutation combined with upregulated TP63 expression linked to tumorigenesis and metastasis development.<br>*Prognosis:* 13.3 survival months | Transcriptional networks linked to later stages of pancreatic development and differentiation. Upregulated transcription factors (NR5A2, MIST1 and RBPJL) involved in acinar cell differentiation and regeneration after pancreatitis. Activated genes linked to endocrine differentiation and MODY diabetes, Exocrine secretion and regulation of beta cell development.<br>*Prognosis:* 25.6 survival months | B and T cells infiltration. Expressed genes involved in antigen presentation and in B cell, CD4+ T cell, CD8+ T cell and Toll-like receptor signaling pathways. CTLA4 and PD1 upregulated and linked to immune suppression.<br>*Prognosis:* 30 survival months |
| Collisson | Classical | Quasi-Mesenchymal | Exocrine-like | - |
|  | High epithelial and cell adhesion-associated (GATA6) gene expression. KRAS mutation dependent.<br>*Prognosis:* 23 months survival | Upregulation of Mesenchyme associated genes<br>*Prognosis:* 6.6 months survival | Upregulated Tumor digestive exocrine enzyme genes.<br>*Prognosis:* 19.7 months survival |  |
| Moffitt | Classical | Basal | - | - |
|  | Overexpressed adhesion-associated, ribosomal and epithelial genes (GATA6).<br>*Prognosis:* 19 months survival | Overexpressed mesenchymal genes, also known to be upregulated in the Basal subtype of breast and bladder cancer.<br>*Prognosis:* 11 months survival |  |  |

### State of the art for gene signatures validation and subtypes reproducibility

More than one study raised skepticism about the generalizability of the subtypes established by Collisson *et al*., Moffitt *et al*., and Bailey *et al*. found discrepancies in the biological relevance of the tumor classes, motivating us and others to reassess these findings through additional analyses.

Birnbaum *et al*. [15] assessed the prognostic value of the three published classifiers by using a large cohort of primary tumor samples coming from 15 different public datasets. Their results independently confirmed the prognostic value of the Moffitt *et al*. and Bailey *et al*. gene classifiers set but not the Collisson *et al*. classifiers.

The Cancer Genome Atlas (TCGA) Research Network [16] performed the same unsupervised analysis as Collisson *et al*., Moffitt *et al*. and Bailey *et al*. on their cohort of primary tumor samples, using each time the corresponding list of gene signatures. Once the clusters had been identified, they investigated their relationship with the purity of the tumor samples. They reported that subtypes such as Immunogenic, ADEX (from Bailey *et al*.) and Exocrine-like (from Collisson *et al*.) are related to low tumor purity, suggesting a contamination of the tumor samples from adjacent normal pancreatic tissue.

Rashid *et al*. [17] explored the ability of the three proposed gene set classifiers in prognostic differentiation and subtype replicability. They applied consensus clustering on nine independent patient cohorts for assessing clustering robustness and carried out survival analysis on those cohorts where survival information was available. They noticed that the two tumor-specific subtypes presented by Moffitt *et al*. are consistent across datasets and thus robust in reproducing different patient subgroups that are prognostically relevant.

Janky *et al*. [18], used the 62 genes proposed by Collisson *et al*. on a larger cohort of whole tumor samples. The three groups obtained from clustering the data showed an almost perfect overlap with the existing subtypes. However, the prognosis related to such clusters appears to be partially inconsistent with respect to the original study. Collisson *et al*. associated the Exocrine-like subtype with good prognosis, contrary to Janky *et al*. that found the subtype associated with bad prognosis, denoting inability of the signature to distinguish the subtypes according to the survival outcome assigned by the study.

### Inconsistencies of the subtypes

Collisson *et al*. themselves recognized that the Exocrine-like subtype might be the consequence of the presence of normal tissue exocrine cells in the sample. This was noticed while observing the subtypes including *in vitro* data, which were not linked to the Exocrine-like subtype, confirmed by Moffitt *et al*.. Similarly, several critiques have been directed at the ADEX subtype of Bailey *et al*., which shows discordance in the subtype assignment and suggests a contamination of acinar cells [16,19].

The existence of the immunogenic subtype of Bailey *et al*. has also been subject to doubts. Maurer *et al*. [20] noticed that samples classified as Immunogenic showed an enrichment in the stromal compartment, which might explain that Immunogenic subtype originates from the tumor microenvironment. Sivakumar *et al*. [21] determined the functional pathways involved in each of the Bailey *et al*. subtypes and found a link between the Immunogenic subtype and the cell cycle signaling pathway. Their samples appear to be low immunogenic.

However, to verify that its discovery was not a direct consequence of dense immunological samples, its composition requires additional examination.

### Problem statement

A key observation is that the molecular subtypes deduced by the three studies are based on independent cohorts of patients with very diverse sample characteristics: Collisson *et al*. partially used laser microdissected samples, contrary to Moffitt *et al*. who microdissected the tumor from the stroma compartment virtually (*in silico*). Bailey *et al*. considered only samples with high epithelial content (>40%) whose histological subtype is not only ductal, while Collisson *et al*. and Moffitt *et al*. focused their attention on the ductal subtype alone. Data with different properties might induce inconsistent results in a comparison. Specifically, the different subtypes may simply reflect differences in the composition of the samples and hence the generalization and meaning of related biomarkers on different datasets remains unclear.

Most of the existing validation studies assess the prognostic power of the signatures by survival analysis on the patient clusters. Several studies investigated the composition of the public subtypes by observing the tumor compartment and the cellular environment of the samples. Nevertheless, a systematic study assessing the robustness of the signatures across different datasets is lacking. Here we seek to answer the question if it is possible to replicate the same subgroups in a cohort that is not the one used for signature discovery. Moreover, it has been shown previously in breast cancer that random gene signatures often show performance comparable to published signatures [22]. Hence, we also compare the performance of published signatures to randomly generated signatures of the same size (w.r.t. number of marker genes). In addition to survival analysis, which has been investigated before, we also link PDAC signatures to differences in cell type composition and compare them w.r.t. functional enrichment. With the latter we shed light on the sample characteristics that the PDAC signatures represent. We demonstrate that they are only indirectly, if it all, linked to cancer subtypes.

## MATERIALS AND METHODS

### Statistical analysis

Strictly standardized mean difference (SSMD) was used to assess the difference in expression of the signatures between the tumor subgroups as robustly as possible. Taking each gene in the signature and the three dataset used for their discovery, we computed pairwise SSMD between subtypes and saved the SSMD that is highest in absolute value. The distribution of highest SSMD for each signature in each dataset is shown as a density curve. Gene expression distributions of the subtypes for each gene in the signature are also compared with a two-sided Wilcoxon test and Kruskal-Wallis test (Bonferroni corrected p-values <0.05) when comparing two or more groups, respectively.

### Clustering

Agglomerative hierarchical clustering was implemented with the SciPy 1.6.0 Python library with correlation as metric. The analysis was applied to each dataset to identify signatures-related clusters, after filtering the gene expression matrix for signature genes. All data were z-score transformed before clustering. The number of examined clusters were two, three or four depending on the expected number of subtypes according to Moffitt *et al*., Collisson *et al*. or Bailey *et al*., respectively. We investigated the cluster differences using on each gene in the signature a two-sided Wilcoxon rank sum test for two and Kruskal-Wallis test for three and four clusters and taking the Bonferroni adjusted p-value (<0.05) distribution.

More details about the signatures overlapping which each datasets can be found in Supplementary Table S2.

### Classification

Datasets used for classification were batch-corrected in order to avoid confounding between different sample preparation and origin. Correction was performed with the RemoveBatchEffect function included in limma 3.48.1 R package. Supplementary Figure S1 shows a principal component analysis of the eight datasets before and after correction. Only genes in common across all the dataset were considered for supervised analysis (more details in Supplementary table S3). Data were z-score transformed before classification.

Classification was performed with the Random Forest Classifier in the Python package Scikit-learn 0.24.2, keeping the default values as parameters. Nine prediction models were built employing the three molecular signatures on the three datasets used for their discovery. Considering one dataset at a time, we filter the genes of each signature to employ them as predictive features. Samples in the current training dataset have available subtype labels which are used as target variables for the model. Each model is applied to a set of eight test datasets where samples are classified according to the target variable. The nine models are built with the aim of classifying different datasets into the three subtype schemes by using the three signatures. A graphic illustration of the model construction is shown in Figure 2. Average accuracy from 5-fold cross validation is used as an evaluation metric.

**Figure 1.**
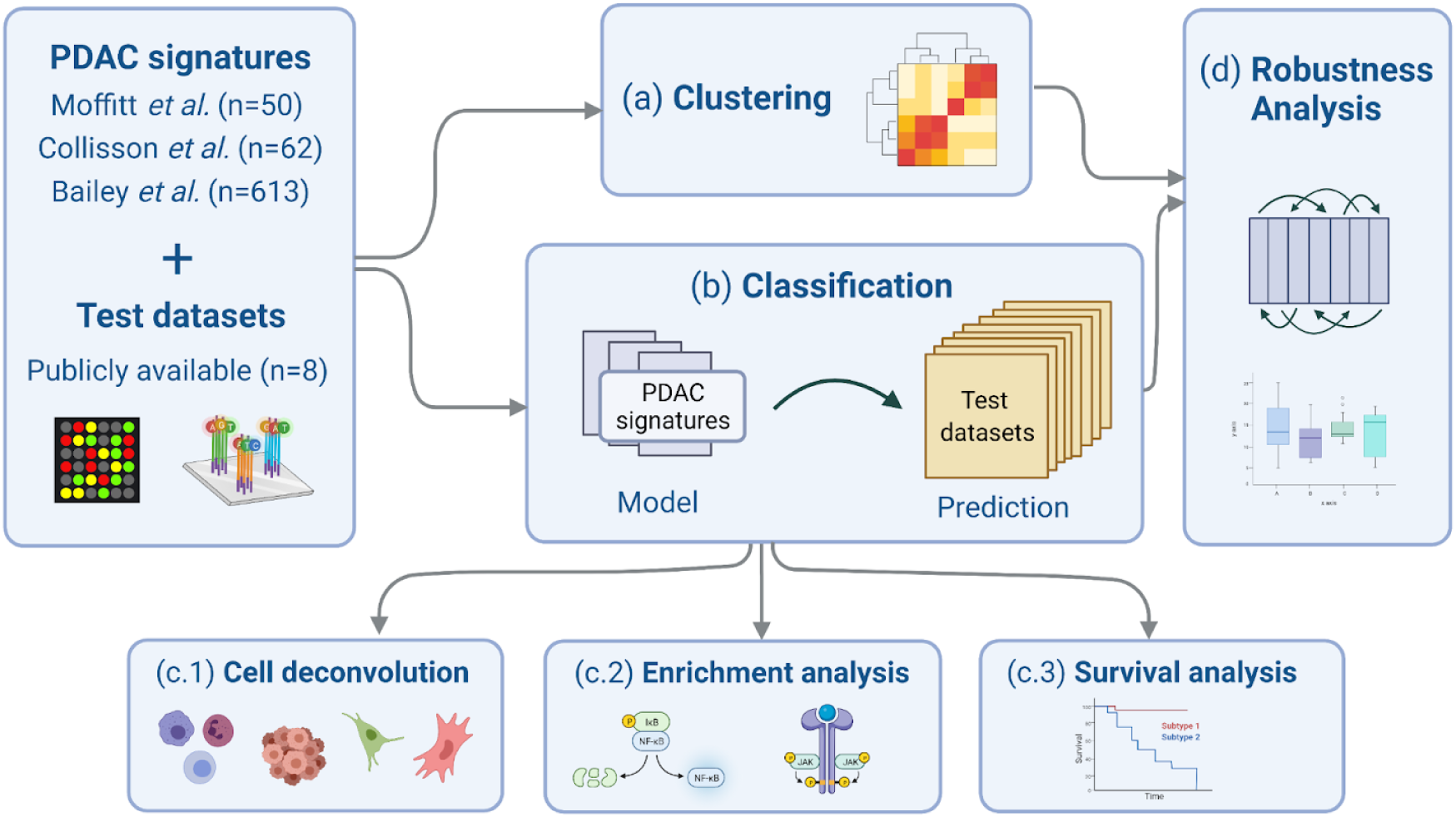
After collecting the PDAC signatures and the eight test datasets, we perform (a) clustering and (b) classification analysis. Based on the latter, we follow up with compositional analysis of the predicted subtypes *via* (c.1) *in silico* cell type deconvolution, (c.2) functional enrichment analysis, and (c.3) survival analysis. (d) Finally, we also perform robustness analysis to learn if random signatures have comparable predictive power.

**Figure 2.**
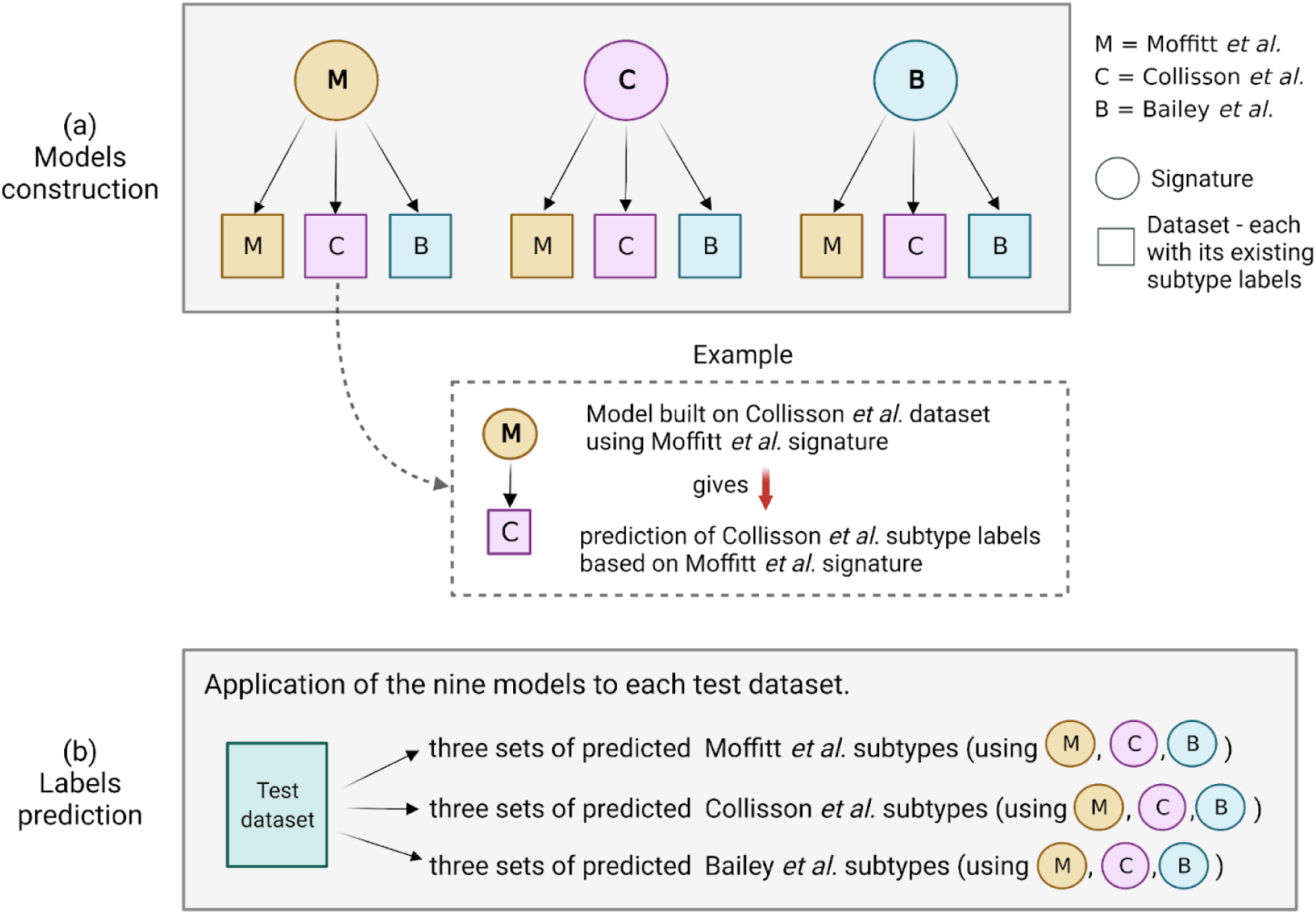
Illustration of the prediction models framework. (a) Models are trained using the three available signatures as a set of predictive features and changing, in turn, the training dataset, considering the one used by Moffitt *et al*., Collisson *et al*. and Bailey *et al*.. This results in nine models that are applied to eight test datasets. (b) Each model predicts the subtype labels linked to the training dataset: if we consider, for instance, a model trained on Moffitt *et al*. dataset, each test dataset will have three sets of predicted Moffitt *et al*. subtypes, obtained using the three signatures.

Besides their predictive power, we want to assess whether the signatures stratify PDAC samples in subgroups which correspond to the subtypes the authors outlined. To this end, we investigated the predicted subtypes in their cell environment composition, their functional mechanisms and their survival differences.

### Robustness analysis

Robustness of the signatures was assessed comparing their performance to random gene sets of the same size. Comparison is carried out using both clustering and classification. In classification, we run the models with three settings. First using as features the real signatures, then the random signatures and, lastly, shuffling the subtype labels while keeping the real signatures. We performed 1000 runs for each setting. Each average of the 5-fold cross-validation accuracy was stored and used for evaluation.We selected the signatures from every dataset and used them to generate clusters. Random signatures were used for the same purpose. The similarity between clusters obtained using the real signatures and random signatures are compared by computing the Adjusted Rand Index (ARI). A pairwise ARI was used to compare clusters obtained from the random signatures. SSMD was used to assess the difference between ARI of the real signatures *vs*. random signatures-derived clusters and the pairwise ARI computed on random signatures-derived clusters. These analyses were performed in Python with Scikit-learn 0.24.2.

### Cell deconvolution and ssGSEA

We estimated the cellular microenvironment by deconvoluting bulk RNAseq data with the CIBERSORTx [23] method. Human pancreas single-cell transcriptome was obtained from Tosti *et al*. [24] and used to obtain a cell-type specific signature matrix for the deconvolution tool.

Single sample gene set enrichment analysis (ssGSEA) was performed in Python using the package GSEApy 0.10.5 [25],[26],[27]. KEGG [28], Reactome [29] and Gene Ontology terms (biological processes, molecular functions, cellular components) [30]. ssGSEA can be understood as a gene-set level aggregated score that reflects a pathway or gene set’s activity where genes in the set are up- or down-regulated in a coordinated, i.e. non-random fashion. Enrichment scores thus allow comparing pathway activity across samples and we use ANOVA and two-sided t-test to identify enriched terms that reflect the phenotypic differences across the predicted subtypes from a functional pathways perspective (Bonferroni corrected p-values <0.05). The term with the highest mean enrichment score is marked as enriched in that subtype. The summary tables in the supplement show, for each enrichment library, the top 30 terms found significantly enriched in most of the cohorts and significantly associated with the different classes predicted by the nine classification models. For each subtype, we define a term upregulated or downregulated in that subtype by looking at the sign of the corresponding enrichment score.

### Survival analysis

Survival analysis was conducted with the Python package Lifelines 0.25.9 for datasets where survival information was available. Survival difference was assessed by computing pairwise log-rank tests between groups.

### Software and tools

Analyses were performed in Python 3.7.3 and R 4.1.0.

Scripts to reproduce analyses and figures are available on GitHub at https://github.com/biomedbigdata/PDAC-molecular-classifier-validation.

## RESULTS

### Statistical analysis

We collected RNAseq and array expression profiles from eight datasets publicly available, including the data used by Moffitt *et al*., Collisson *et al*. and Bailey *et al*. as discovery cohorts. For subtypes to be reliably discriminated, signature gene expression values should vary between tumor subgroups. We quantified the expression effect size of each individual gene in the signature by looking at the SSMD across the subtypes of the three studies. As shown in Figure 3 (A), the signature proposed by Bailey *et al*. recorded the highest SSMD in all the three dataset, meaning the majority of the genes in the signature can catch differences between subtypes better than the other two signatures. In contrast, the SSMD distribution obtained using Moffitt *et al*. signature shows the lowest values. Nevertheless, SSMD distributions differ by dataset even when using the same signature. Next, we tested for each individual gene if it is significantly associated with the subtypes (Wilcoxon rank sum test for two subtypes, Kruskal-Wallis test otherwise), as we hypothesize that signatures should be rich in genes that differ significantly across subtypes (Figure 3 (B)).

**Figure 3.**
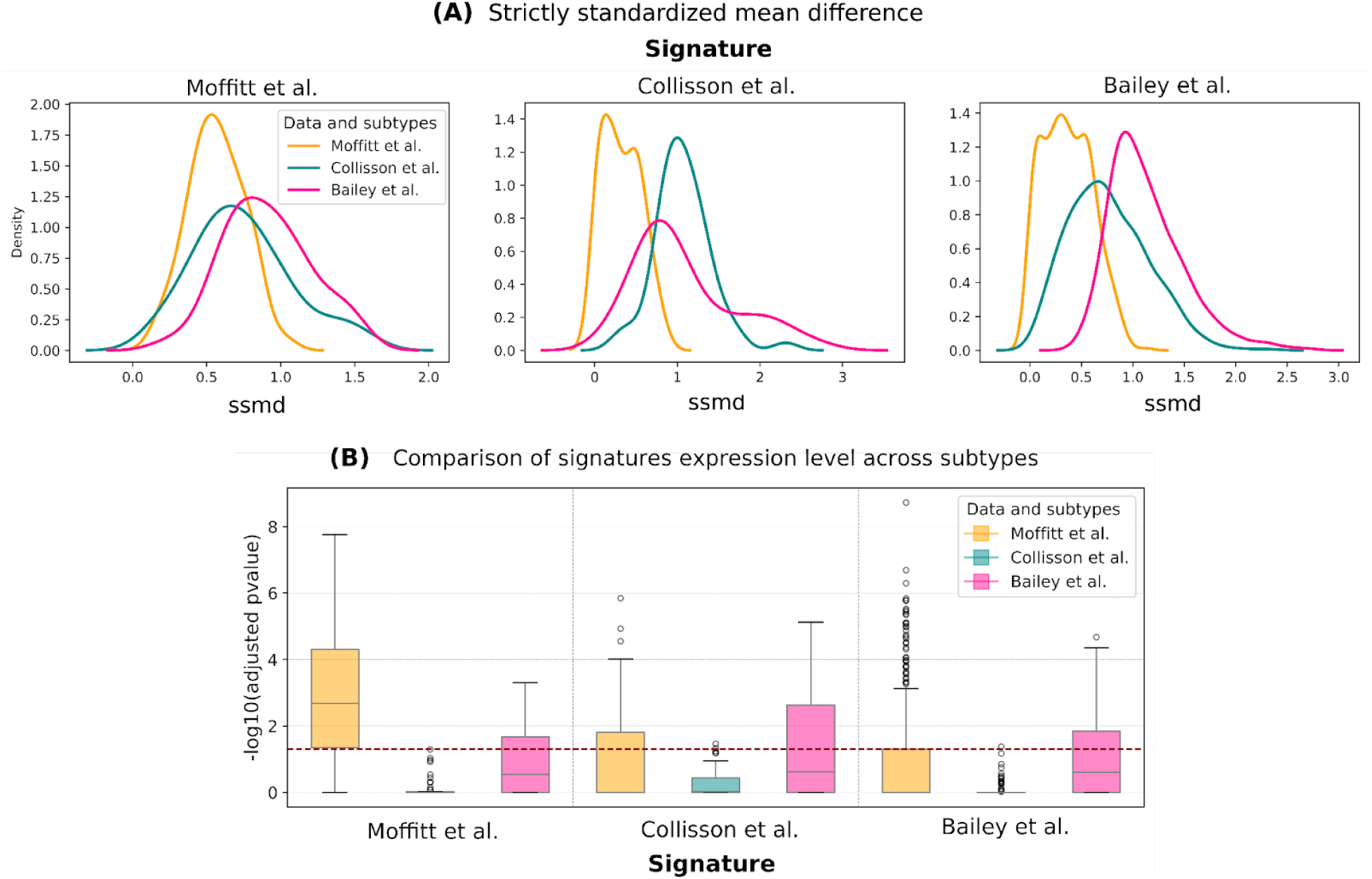
(A) strictly standardized mean difference (SSMD) of the signatures from Moffitt *et al*., Collisson *et al*. and Bailey *et al*. in their discovery datasets. The figure, divided by signatures, shows their effect size in the datasets employed by the three studies and highlights the weakest SSMD characterizing Moffitt *et al*. signatures when compared to the other two. (B) for each signature we assessed its variability across subtypes. Considering one gene at a time, we computed Wilcoxon rank sum test for two-class comparison and Kruskal-Wallis for the three and four-class. The p-values returned by each test were adjusted with Bonferroni correction and are shown log-transformed. The horizontal dashed line indicates the threshold of significance (p-value=0.05). The x-axis indicates the different signatures while the color indicates different subtypes.

We further examined the consistency of the signatures across independent datasets in unsupervised and supervised analysis, followed by a comparison against random gene signatures and a functional evaluation of the subtypes.

### Clustering

Each of the eight datasets was subject to hierarchical clustering using only signature genes. We would expect that the resulting clusters reflect subtypes across different datasets. However, we observed that signature-based clusters accurately reflect subtypes only in their discovery dataset (Figure 4 (A.1), (B.2) and (C.3)). Although we found an overlap between some of the clusters and existing labels, we decided to quantify and assess the difference in expression between clusters. To do so, we used a Wilcoxon rank sum test for two and Kruskal-Wallis test for three and four clusters, and showed the adjusted p-value distribution (Figure 4 below every heatmap).

**Figure 4.**
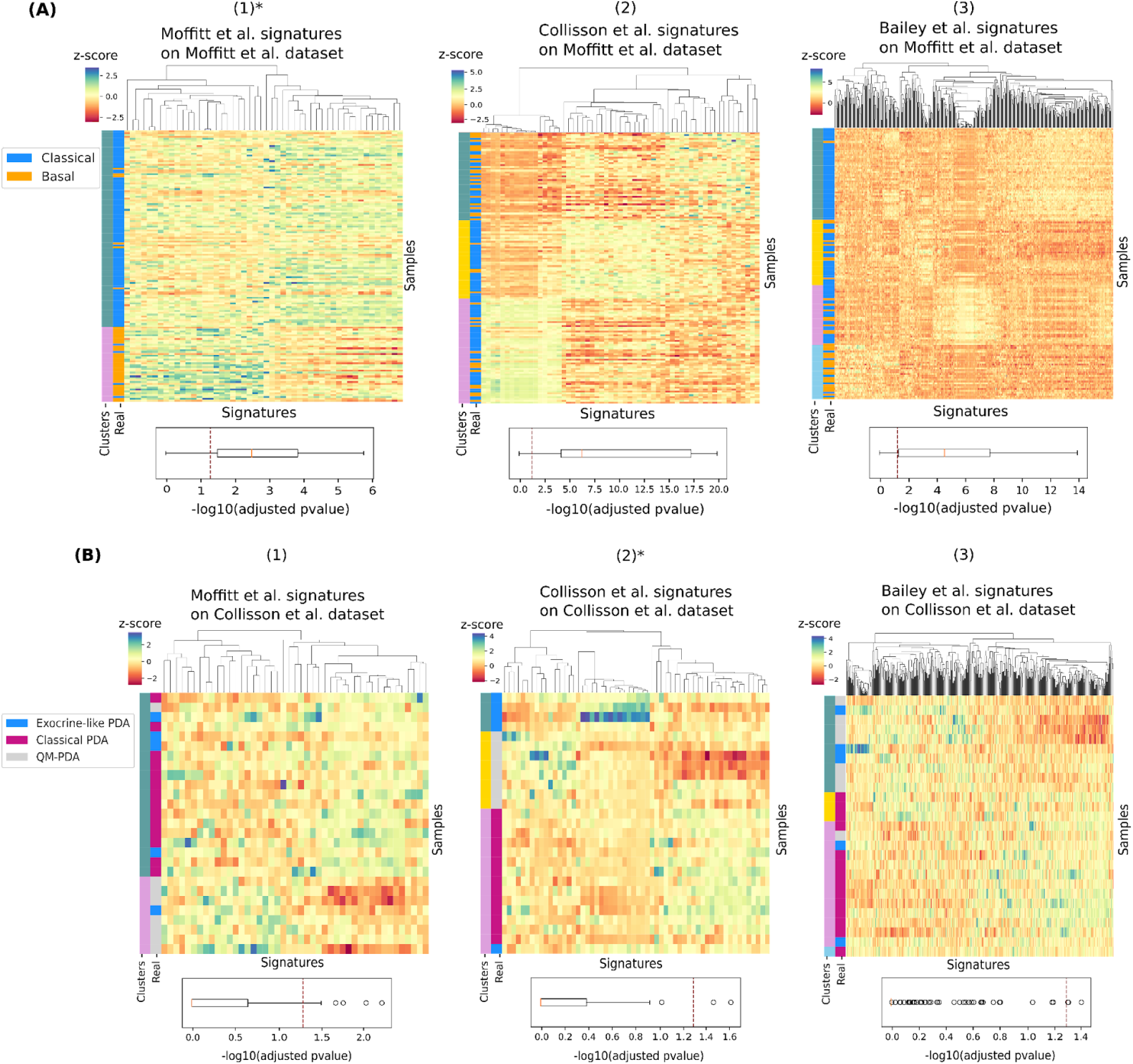
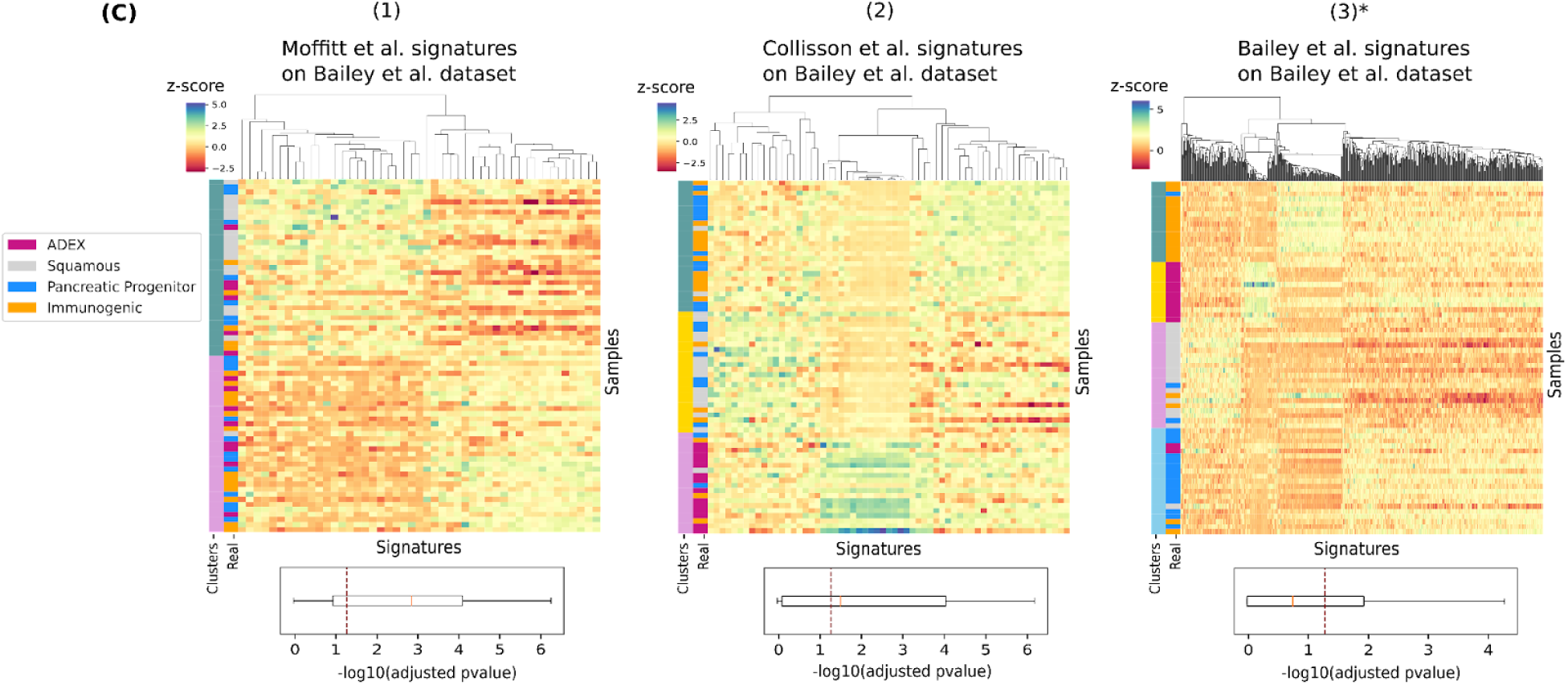
Hierarchical clustering of the Moffitt *et al*. (A), Collisson *et al*. (B) and Bailey *et al*. (C) dataset, performed using Moffitt *et al*. (1), Collisson *et al*. (2) and Bailey *et al*. (2) signatures. Overlap between predicted clusters and real subtypes is observed on the left side of each heatmap, where samples are sorted by clusters. For each gene, association between signature expression profile and clusters is displayed with -log10(adjusted p-values) distribution below every heatmap, where p-values were obtained with Wilcoxon rank sum test for two and Kruskal-Wallis test for three and four clusters. Significance threshold of p-value=0.05 is indicated with a vertical dashed line. (*) Signatures applied on the same dataset used for their discovery.

Considering the Collisson *et al*. dataset, we observed a perfect overlap between existing Collisson *et al*. subtype labels and identified clusters on its discovery dataset (Figure 4 (B.2)) and a good overlap for the Moffitt *et al*. subtypes (Figure 4 (B.1)). However, the expression profiles of the gene signatures do not seem to discriminate well between the clusters, as indicated by the -log10(adjusted p-value) distribution. The same outcome was observed in Figure 4 (C.3), where the Bailey *et al*. signature perfectly clusters the samples into their assigned subtypes while showing high similarity in expression levels between clusters. Clustering of other datasets is available in Supplementary Figure S2.

### Classification

A limitation when comparing subtypes across different studies is that the corresponding subtype labels are only available for the discovery cohort but not for the others. To enable further analysis and to assess the predictive potential of each signature, we build classification models that allow us to predict subtype labels for any of the three subtype schemes. Specifically, we use the Moffitt *et al*., Collisson *et al*. and Bailey *et al*. datasets for training a random forest subtype classifier. For each of the different signatures, we build three models using the three datasets and their existing subtype labels, respectively, as illustrated in Figure 2. We use the nine models to classify each of the test datasets. When comparing predicted and original labels (for those datasets where PDAC molecular subtype labels were available), we observe that the real labels are mixed among the predicted labels (Figure S3). This suggests that predictions are either not robust or that the different subtype schemes do not agree very well, as also suggested by the clustering analysis.

To test if predictions are robust, we calculated the mean accuracy in a 1000 times repeated K-fold cross-validation where, in addition to the Moffitt *et al*., Collisson *et al*. and Bailey *et al*. datasets, we also included the Badea *et al*. dataset, which comes with already assigned Collisson *et al*. tumor subtype labels. As shown in Figure 5 the Moffitt *et al*. signature showed the most dramatic differences in prediction accuracy. All the three signatures gave the best performance on the Moffitt *et al*. dataset for the prediction of their subtypes (Basal and Classical). Such high accuracy is likely owing to the simpler binary classification task. The Bailey *et al*. signature, with the largest number of genes, performs best across all datasets.

**Figure 5.**
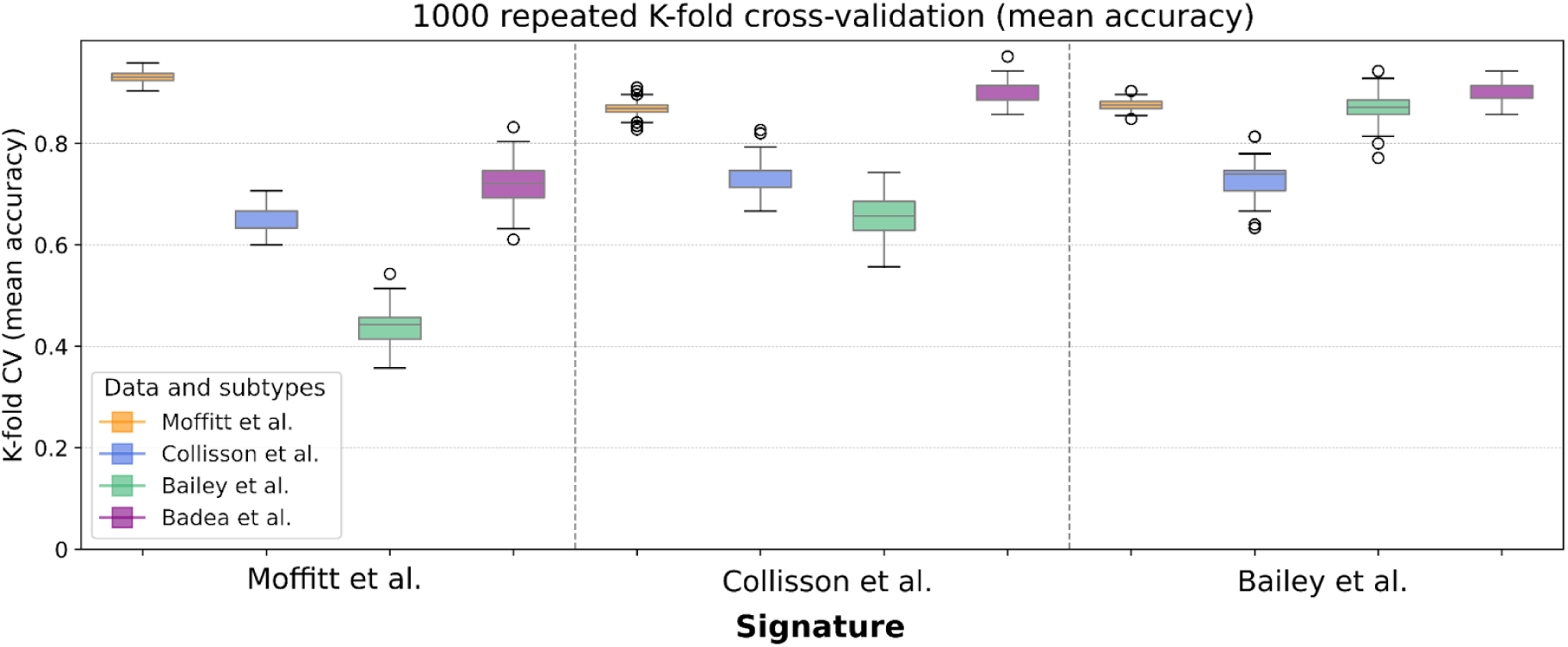
Distribution of the mean accuracy obtained by 1000 repeated K-fold cross validation of classification. Use of signature from Moffitt *et al*. (left), Collisson *et al*. (center) and Bailey *et al*. (right) on the four datasets used for model training. Models built on Moffitt *et al*. mixed tissue dataset show better performance when compared with models built on exclusive tumor data.

### Robustness analysis

Several studies have shown that random gene signatures can outperform literature-reported signatures in outcome prediction [22,31], underlining that signature genes do not necessarily capture tumor biology. We hypothesize that random gene signatures may also be able to outperform published PDAC signatures in subtype prediction. To investigate this hypothesis, we took the models trained on the Moffitt *et al*., Collisson *et al*. and Bailey *et al*. datasets and, for each dataset, compared the performance of a baseline model using signature genes against 1000 random gene sets of the same size as the corresponding signature. We additionally repeated model training after shuffling the subtype labels 1000 times. To assess if any of the models suffers from overfitting we computed K-fold cross validation.

As shown in Figure 6 some of the random signatures reach an accuracy similar to the literature-reported signatures, in some instances even outperforming them. The model trained on whole tumor data used by Moffitt *et al*. shows the best results performing slightly better than random signatures. The Collisson *et al*. and Bailey *et al*. signature models show relatively poor classification performance.

**Figure 6.**
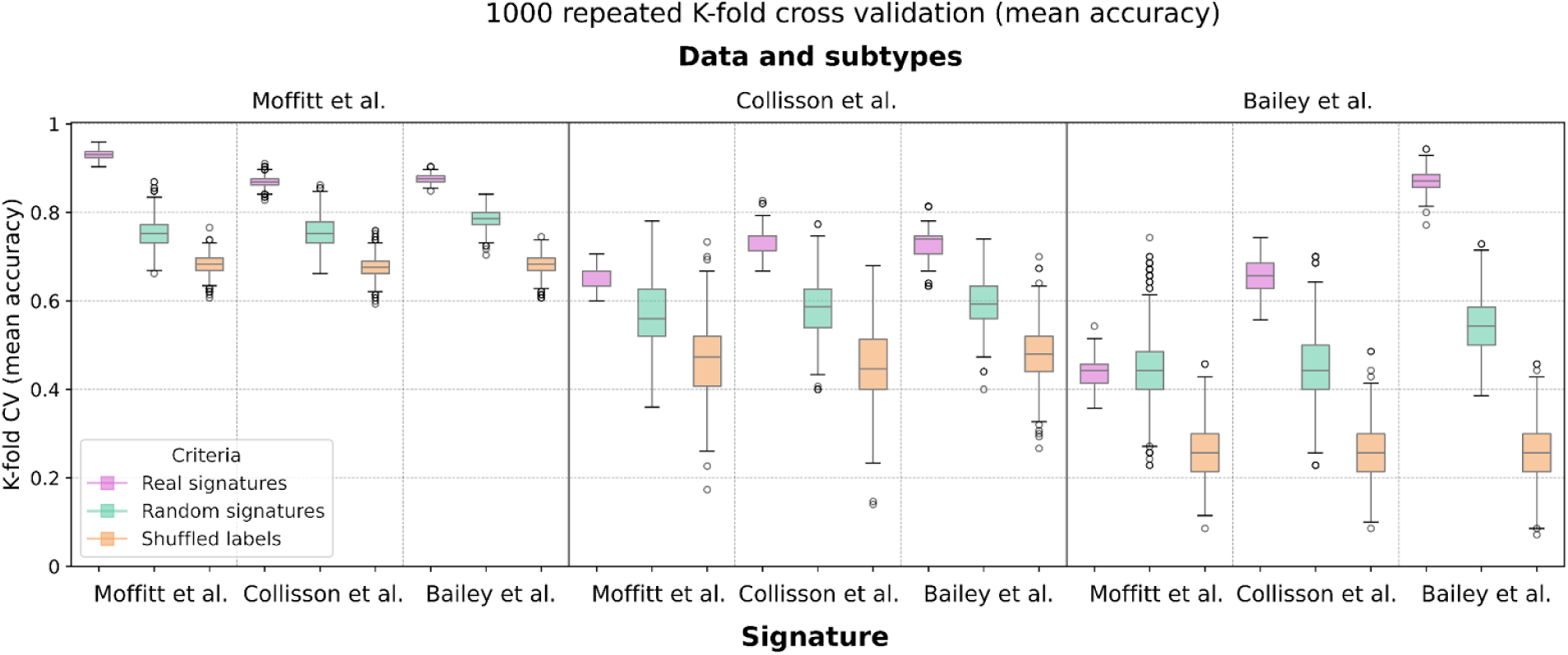
Evaluation of the classification ability of Moffitt *et al*. (left), Collisson *et al*. (center) and Bailey *et al*. (right) signatures used as features for classifying in turn Moffitt *et al*., Collisson *et al*. and Bailey *et al*. datasets. The same task was repeated using random signatures and shuffled labels of the target variable. Models are executed 1000 times with mean accuracy from K-fold cross validation as evaluation metric.

We used the Adjusted Rand Index (ARI) to measure cluster similarity. We expect a low ARI when comparing clusters from random signature and real clusters, ideally much lower than the ARI we obtain when comparing pairwise random against random gene set clusters. Supplementary figure S4 shows that across all eight cohorts most signatures achieve a very low ARI close to zero when compared against random gene set clusters. To summarize these findings, we computed the SSMD [32], which is a robust measure to quantify the difference of the means for the two distributions of ARIs values while accounting for the considerable variance (Figure 7). We notice differences for individual datasets which suggest that the dominant signal is not always related to the subtypes that the signatures aim to recapitulate. Bailey *et al*. signatures registered higher SSMD values with respect to the other two signatures, meaning that their gene panel generates clusters more different than the ones obtained by employing a random list of genes, compared to the other two signatures.

**Figure 7.**
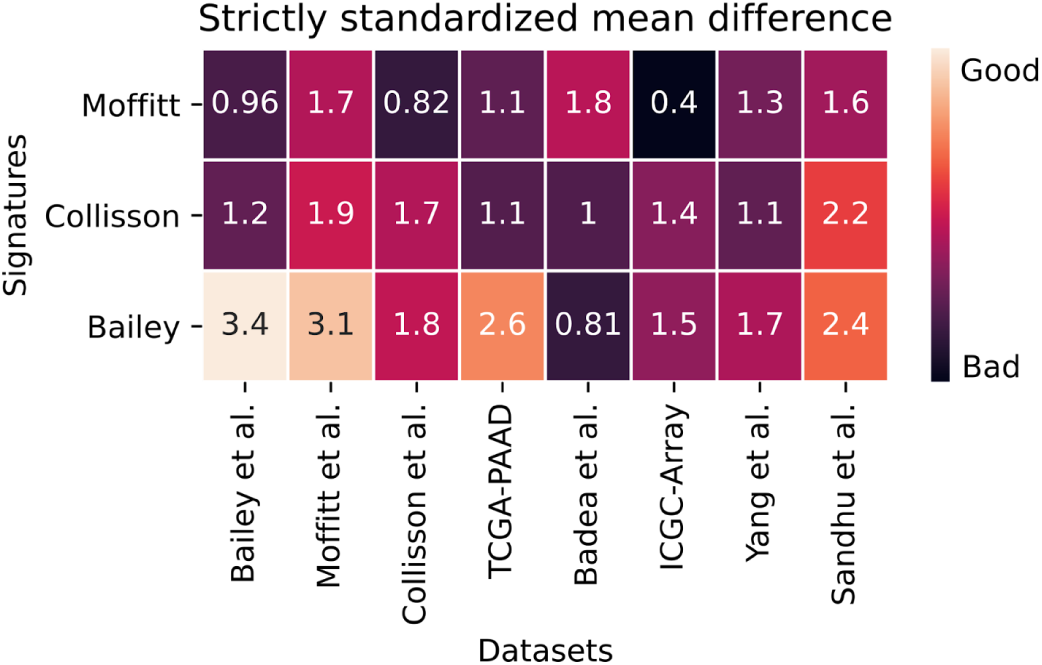
Strictly standardized mean difference (SSMD) comparing two distributions: ARI values describing the overlap of clusters obtained with random signature with real signature-derived clusters are compared to ARI values obtained from pairwised comparison between only random signatures-derived clusters. Large values in the heatmap indicate that signature-derived clusters differ considerably from clusters of random signatures for a given dataset, i.e. the larger the value the better. The values shown are absolute values.

### Cell deconvolution and ssGSEA

With the trained classifiers (see Section Classification), we were able to predict subtype labels, allowing us to investigate the functional characteristics of various subtype definitions across a broad set of datasets. We hypothesize that cellular composition of the samples has a major influence on the bulk transcriptome and that subtype definitions could reflect this heterogeneity rather than properties of the PDAC cells. To investigate if subtype definitions are confounded by stromal or immune cells, we used CIBERSORTx [33] to perform *in silico* cell type deconvolution. Specifically, we used a pancreas single-cell RNA-seq dataset to obtain cell-type-specific signatures allowing us to estimate stromal and immune cell enrichment scores (Supplementary figure S5). As expected, Collisson *et al*. and Bailey *et al*., which use rich tumor samples, are the datasets showing highest enrichment in ductal cells. Notably, we found a high abundance of pancreatic acinar cells in the Bailey *et al*. subtype ADEX and Exocrine-like from Collisson *et al*.. Moffitt subtypes, Basal and Classical, don’t show association to any specific cell type. Subtype schemes show comparable results, not dependent on the signatures but rather dependent on the dataset.

Next, we performed single-sample Gene Set Enrichment Analysis (ssGSEA) to investigate which functional terms are associated with the different subtypes. For each of the nine classification models, we computed sample-wise enrichment scores across functional categories and tested them for significant association with the predicted subtype labels. Pathways we found significantly enriched were mostly linked to metabolic activity, in particular fatty acid and lipid metabolism (Figure 8). Drug metabolism pathways like cytochrome P450 were associated with the Exocrine-like subtype, which is associated with drug resistance which was also observed by Noll *et al. [34]*. (Other ssGSEA summary tables in Supplementary figure S6)

**Figure 8.**
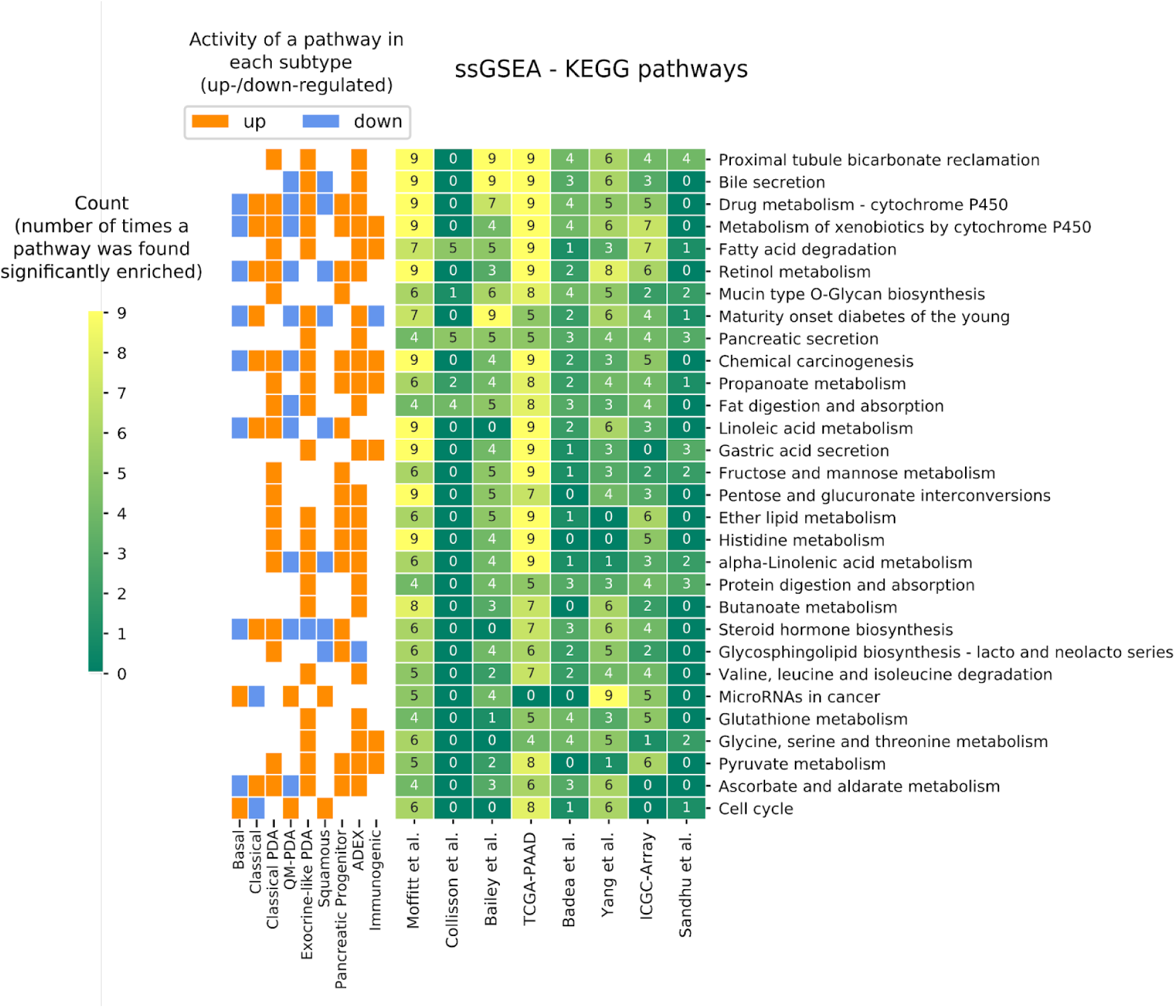
ssGSEA based on KEGG pathways. The table contains the number of times an enrichment term (on the rows) was found significantly enriched in the subtypes comparison for each cohort (on the columns) using the subtypes predicted. Each term can be found enriched up to nine times considering the predictions executed using the nine classification models. On the left, the vertical color bars show whether a pathway is up-/down-regulated and in which subtype. Looking at the top 30 terms that differentiate the most between subtypes, we found decreased metabolic activity in the Basal subtype with respect to the Classical, as shown by the left colorbar indicating down-regulation of metabolic pathways in the Basal subtype.

### Survival analysis

In addition to insights into tumor biology, subtype definitions are used for determining clinically relevant differences in prognosis. We carried out a survival analysis comparing the overall survival of predicted subtypes across those datasets for which survival data was available. Figure 9 shows significant survival differences for the two subtypes proposed by Moffitt *et al*., validating their original findings. For the Collisson *et al*. and Bailey *et al*. signatures the pairwise p-values were only significant in few cases.

**Figure 9.**
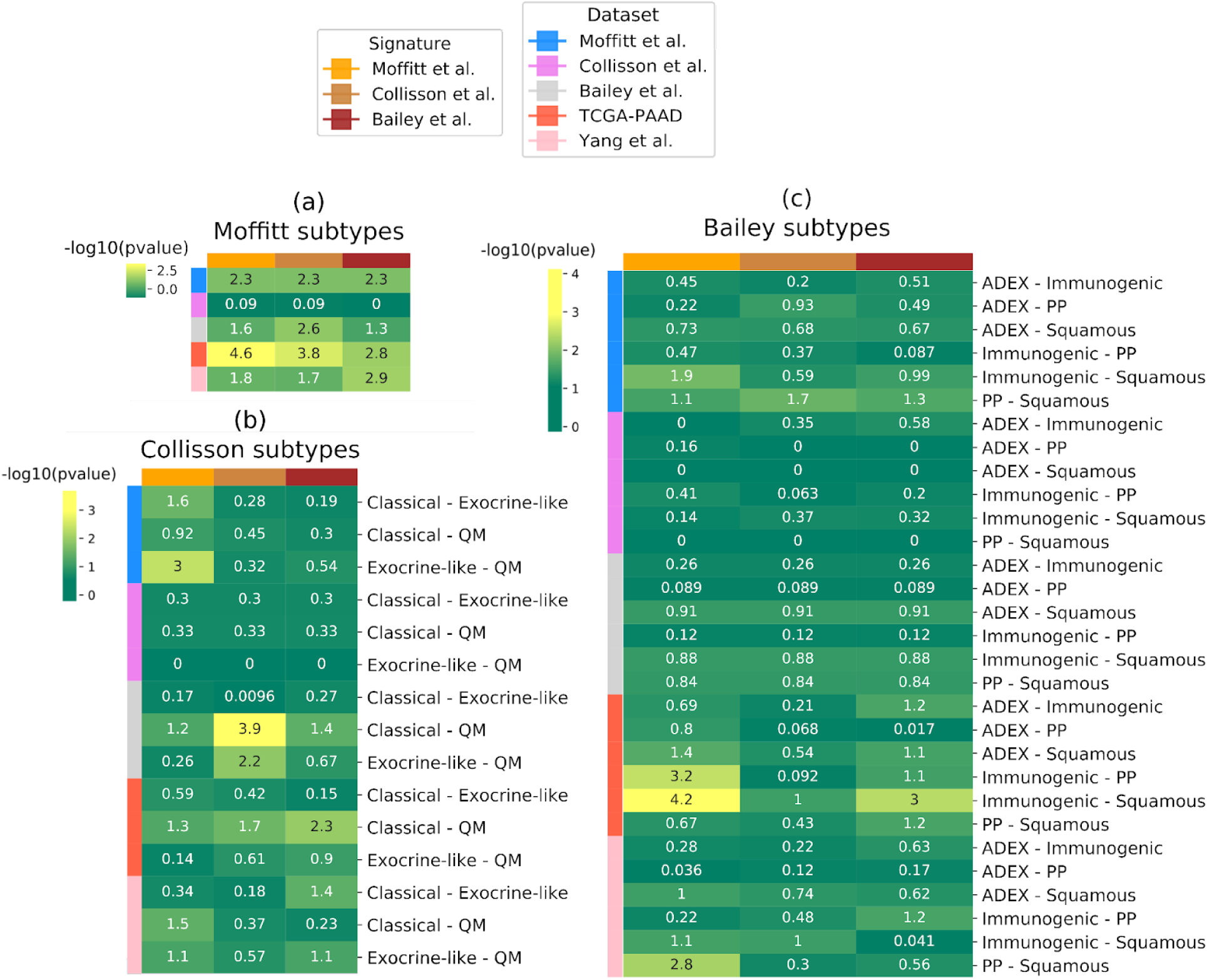
-log10(p-value) from a log-rank test computed comparing overall survival of the predicted subtypes for each dataset whose survival information was available (datasets on the rows). The three subtyping schema (a), (b) and (c) predicted using, in turn, Moffitt *et al*., Collisson *et al*. and Bailey *et al*. signatures. (a) Log-rank test computed between Basal and Classical subtypes from Moffitt *et al*.; (b) pairwise log-rank test between the three Collisson *et al*. subtypes; (c) pairwise log-rank test between the four Bailey *et al*. subtypes.

## DISCUSSION AND CONCLUSION

In this study we evaluated the three major published molecular signatures for PDAC stratification. Our findings indicate that these signatures appear inconsistent when applied to independent datasets, underlining their irreproducibility and further showing how the number and characteristics of PDAC subgroups still remain vague. Our results further suggest that current subtype definitions are mostly driven by tissue composition. In particular, the expression levels of the Bailey *et al*. signature frequently show low association with the four predicted clusters, suggesting that a three subtype schema is more plausible and supporting the hypothesis that one of the four subtypes is an artifact induced by the discovery data used [16,19–21].

Interestingly, the signatures appeared to perform better than random genes, albeit the average classification accuracies are not too far apart, and even random genes achieve good accuracy. Bailey *et al*. signatures performed best in terms of accuracy when compared with the other two signatures along with random signatures or shuffled labels.

In subtype classification, Moffitt *et al*. and Collisson *et al*. signatures (50 and 62 genes, respectively) show less predictive power than Bailey *et al*. signatures (613 genes). The number of features used for a supervised analysis strongly influences the class prediction. Indeed, we observed a difference in accuracy between the first two signatures and the third one, with a plausible link to the amount of features employed for building the predictive model.

Bulk samples deconvoluted at a single-cell resolution revealed how samples enriched of pancreatic acinar cells were classified as Exocrine-like and ADEX, supporting the critique raised by other studies about the authenticity of the two mentioned subtypes, most likely a consequence of normal pancreatic tissue present in the samples that biased the subtype identification [16,18]. Our results present strong evidence that the expression profiles of the three PDAC subtypes along with their signatures are biased by the presence of adjacent normal pancreatic tissue, and suggest that samples with low tumor content were mistaken for an own subtype.

The signatures do not seem to determine prognostically relevant subtypes, as discovered through survival analysis. Indeed, the Basal/Classical subtypes proposed by Moffitt *et al*. are the ones linked to a significantly different outcome, whichever is the signature implemented for classification. Therefore, we hope that future studies will consider the correction or removal of possible confounding factors, such as sample preparation, sampling bias and/or composition. We found the presence of an immunogenic infiltration associated with sample stratification and further focus should be dedicated to the understanding of the immune cell compartment. PDAC seems to generate a strong immune response since the initial state of the disease and it has been proved the impact of tumor-associated macrophages, regulatory T-cells and myeloid-derived suppressor cells on patient prognosis, denoting shorter survival given their prevalence in invasive cancer stages [35–37]. In addition, studies focused on the examination of infiltrating immune-suppressive cells revealed distinct immune cells compartment in distinct tumor groups, suggesting the use of immune response as a strategy for the PDAC stratification [36,38].

Additional assessment can confirm the potential existence of an Immunogenic subtype and bring to light the prognostic role of immune mechanisms. While the work of Mofitt *et al*., Collisson *et al*. and Bailey *et al*. have advanced our understanding of molecular differences within cohorts of PDAC patients, we have to conclude that the proposed signatures are not well suited as a basis for clinical decisions without first accounting for differences in sample preparation and composition. Thus, the use of transcriptome data for PDAC stratification still remains an open challenge. We highlight the importance of an integrated analysis that implements cross-platforms multi-omics data, and we believe that data at a single-cell resolution might provide solid support for an unbiased subtypes definition.

## DATA AVAILABILITY

Eight gene expression datasets were downloaded from public repositories.

Normalized microarray profiles from Moffitt *et al*. and Collisson *et al*. were downloaded from GEO under the accession codes GSE71729 and GSE17891, respectively. Only PDAC primary samples were kept.

Bailey *et al*. RNAseq profiles (ICGC PACA-AU) were obtained by author correspondence in a RSEM counts matrix that we normalized through variance stabilizing transformation in DESeq2. Non-ductal histopathological subtype samples were excluded from this study.

Normalized gene expression array data matrix from ICGC PACA-AU Array, Badea *et al*., Yang *et al*. and Sandhu *et al*. were downloaded from GEO (GSE36924, GSE15471, GSE62452 and GSE60980). Replicates in ICGC PACA-AU array and Badea *et al*. were dropped from this study.

Log2 of the counts for TCGA-PAAD RNAseq were obtained from Xena Browser [39]. After filtering patients with PDAC primary tumor and ductal subtype, data were first converted from logarithmic to counts and then normalized with variance stabilizing transformation in DESeq2.

All probe ids were converted to gene ids, considering the median value if a gene name is mapped to multiple probes. An overview of the datasets used with respective gene and sample size can be found in supplementary table S1.

Molecular signatures and subtype calls assigned to the patients were downloaded from the corresponding publications of Moffitt *et al*., Collisson *et al*. and Bailey *et al*..

## ACKNOWLEDGMENT

Figure 1 and 2 created with BioRender.com.

