## Supplementary Figure for "The Limits of Molecular Signatures for Pancreatic Ductal Adenocarcinoma Subtyping"

### Supplementary Figures

#### Figure S1

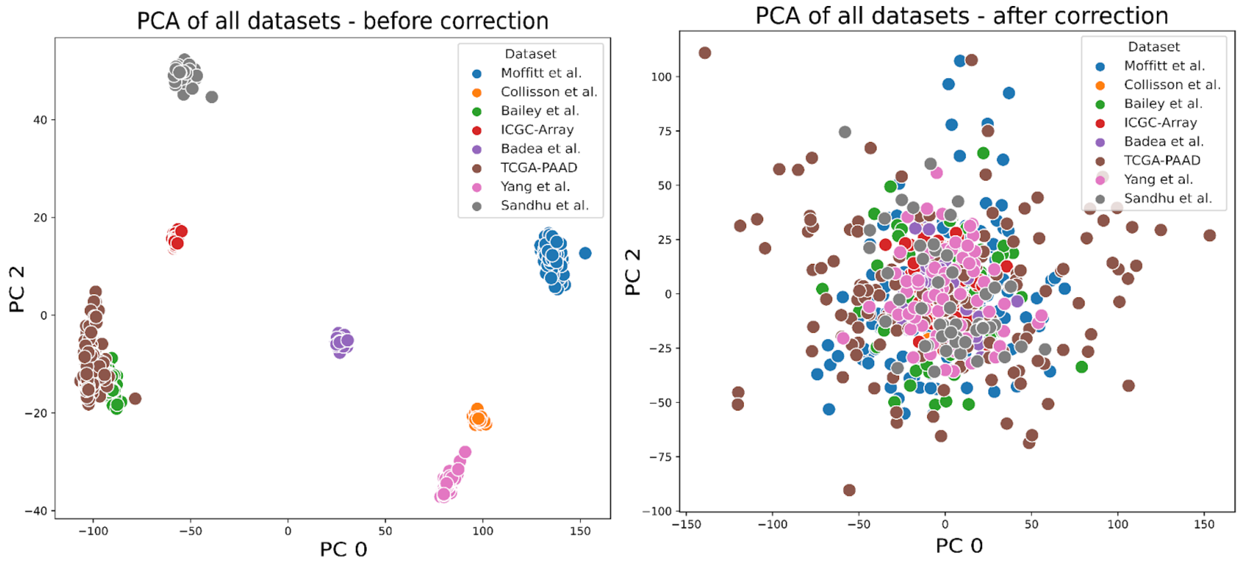


**Figure S1.** Principal Component Analysis (PCA) of the ten validation datasets combined. PCA before correcting the data for different sources of cohorts as a confounding factor (left). The right plot shows the sample of each cohort after the source correction.

#### Figure S2

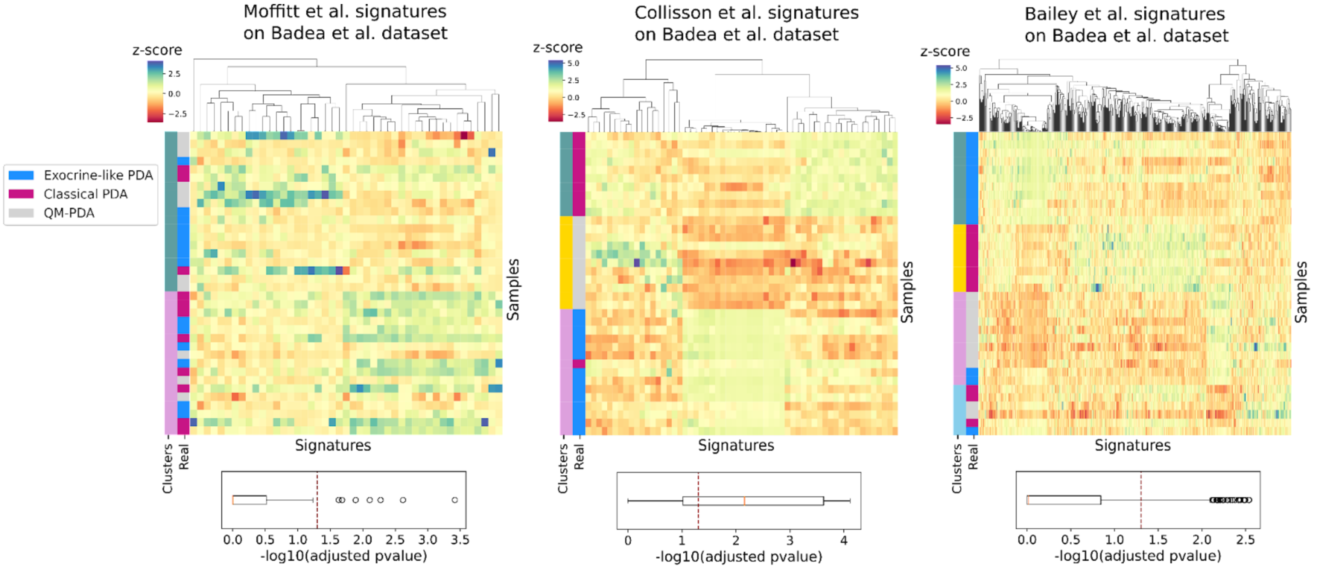

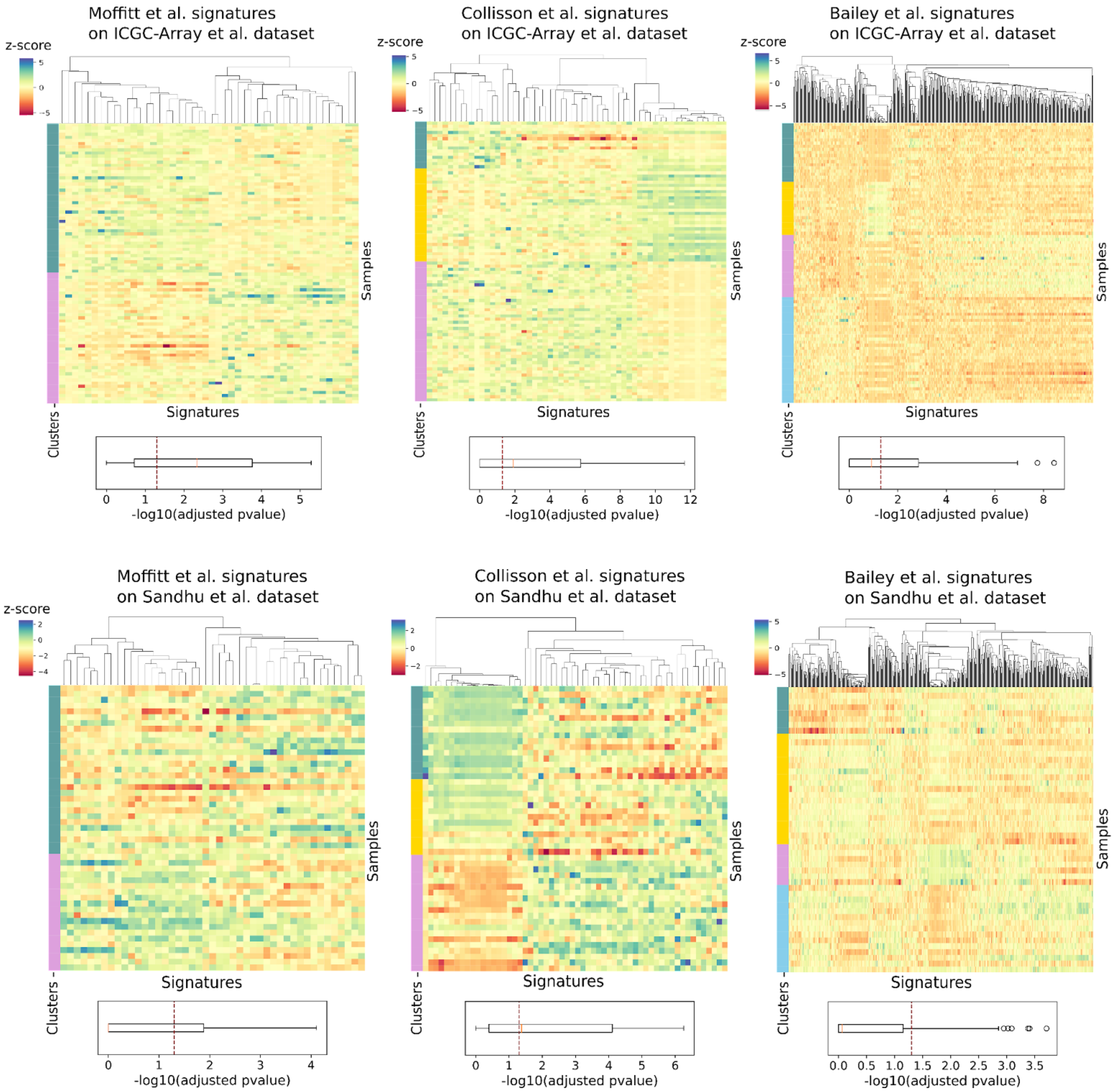
Moffitt et al. signatures
on ICGC-Array et al. dataset
Collisson et al. signatures
on ICGC-Array et al. dataset
Bailey et al. signatures
on ICGC-Array et al. dataset
z-score
Samples
Signatures
Clusters
-log10(adjusted pvalue)
Moffitt et al. signatures
on Sandhu et al. dataset
Collisson et al. signatures
on Sandhu et al. dataset
Bailey et al. signatures
on Sandhu et al. dataset
z-score
Samples
Signatures
Clusters
-log10(adjusted pvalue)

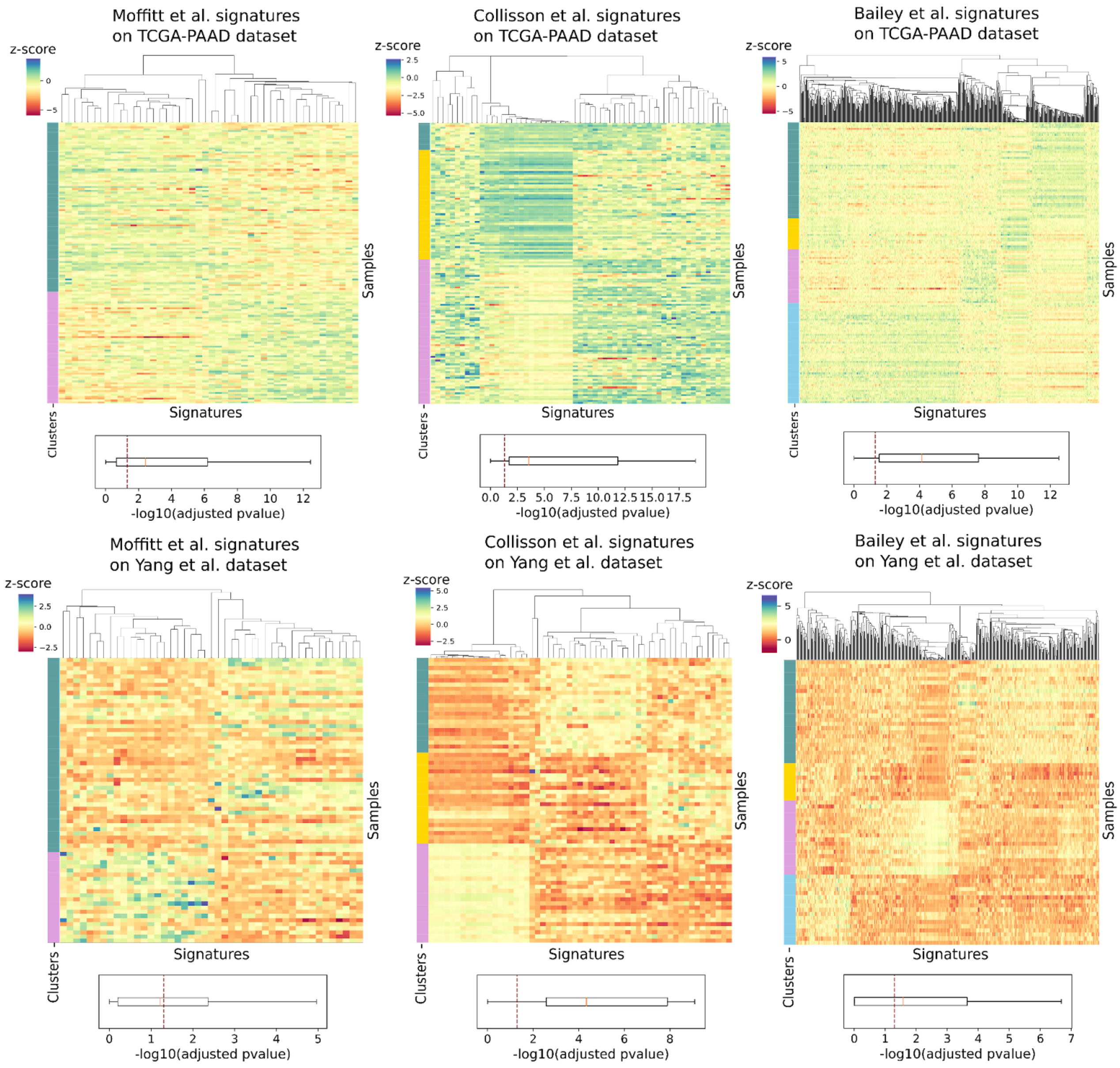


**Figure S2.** Hierarchical clustering of the Badea *et al.*, ICGC-Array, Sandhu *et al.*, TCGA-PAAD, Yang *et al.* z-scored datasets using the signatures from Moffitt *et al.*, Collisson *et al.* and Bailey *et al.*. Below every heatmap, a boxplot shows the -log10(adjusted p-values) distribution obtained assessing the difference in expression between clusters computing, on each gene in the signature, Wilcoxon rank sum test for two and Kruskal-Wallis test for three and four clusters. A vertical dashed line indicates the significance threshold of p-value=0.05.

(*) Clusters of Badea *et al.* are compared with the real subtypes assigned in the study of Collisson *et al.* which returns, as expected, clusters matching the real tumor subgroups.

#### Figure S3

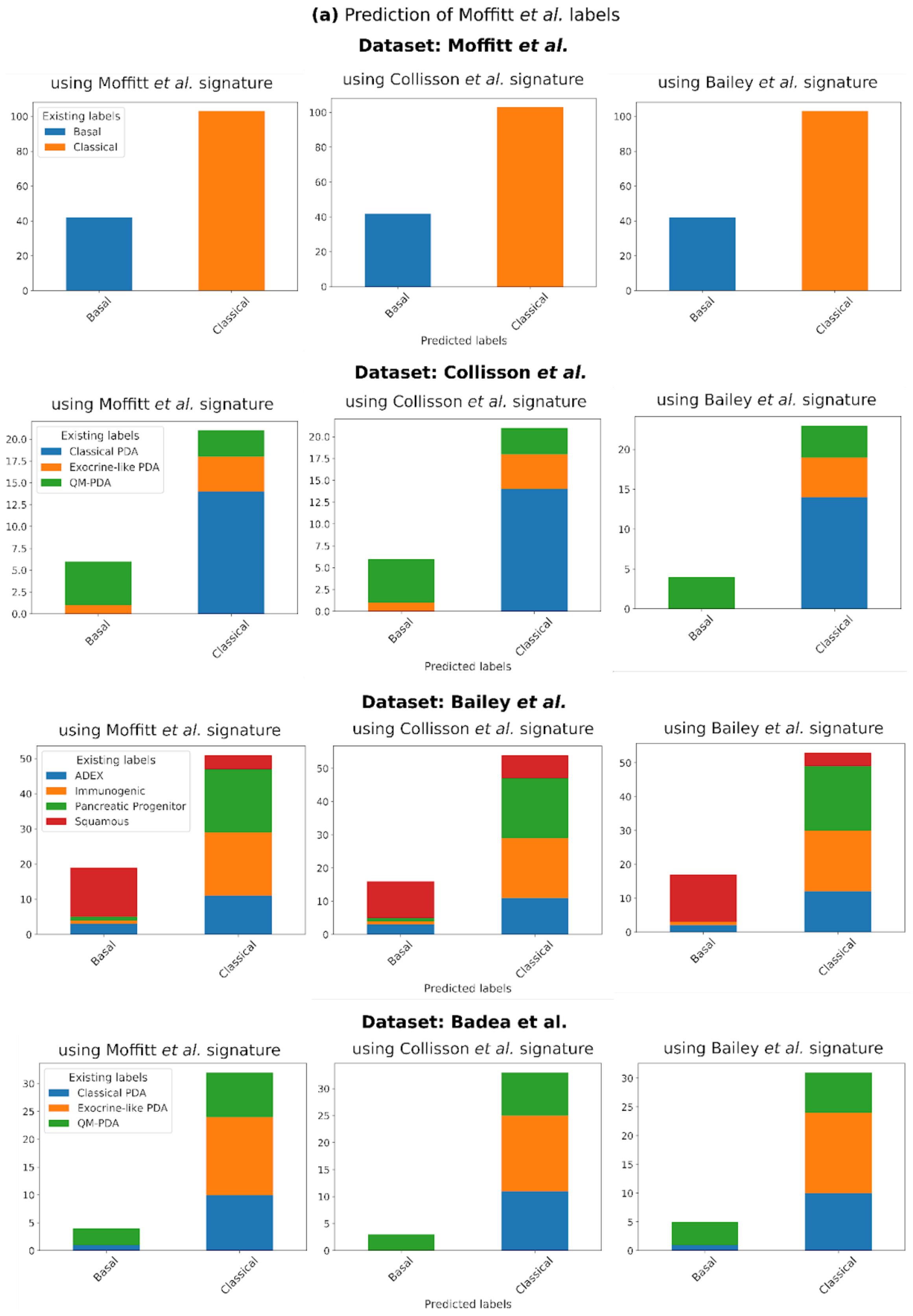

(a) Prediction of Moffitt et al. labels
Dataset: Moffitt et al.
using Moffitt et al. signature
using Collisson et al. signature
using Bailey et al. signature
Existing labels
Basal
Classical
Predicted labels
Dataset: Collisson et al.
Classical PDA
Exocrine-like PDA
QM-PDA
Dataset: Bailey et al.
ADEX
Immunogenic
Pancreatic Progenitor
Squamous
Dataset: Badea et al.

**(b)** Prediction of Collisson *et al.* labels

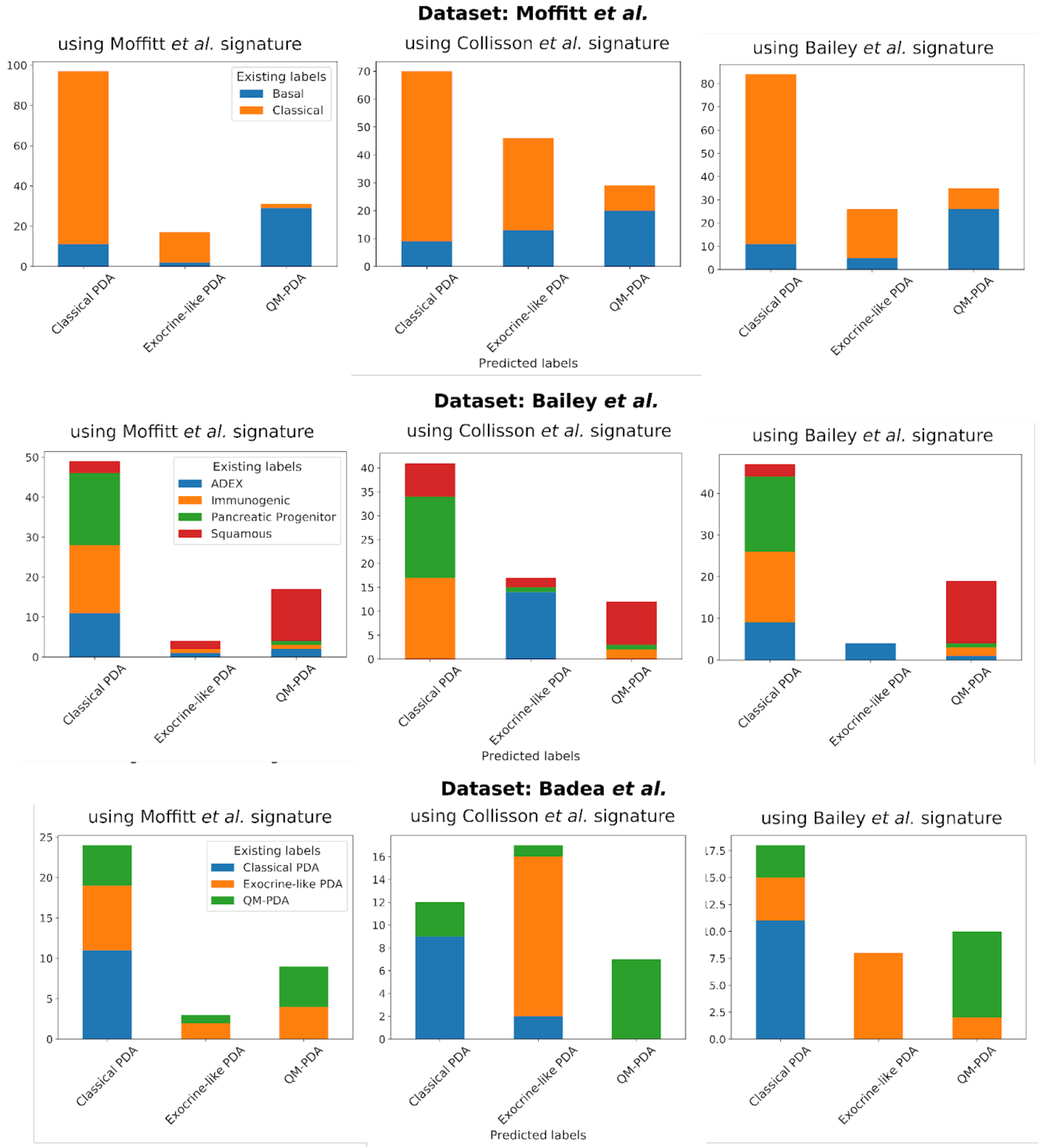

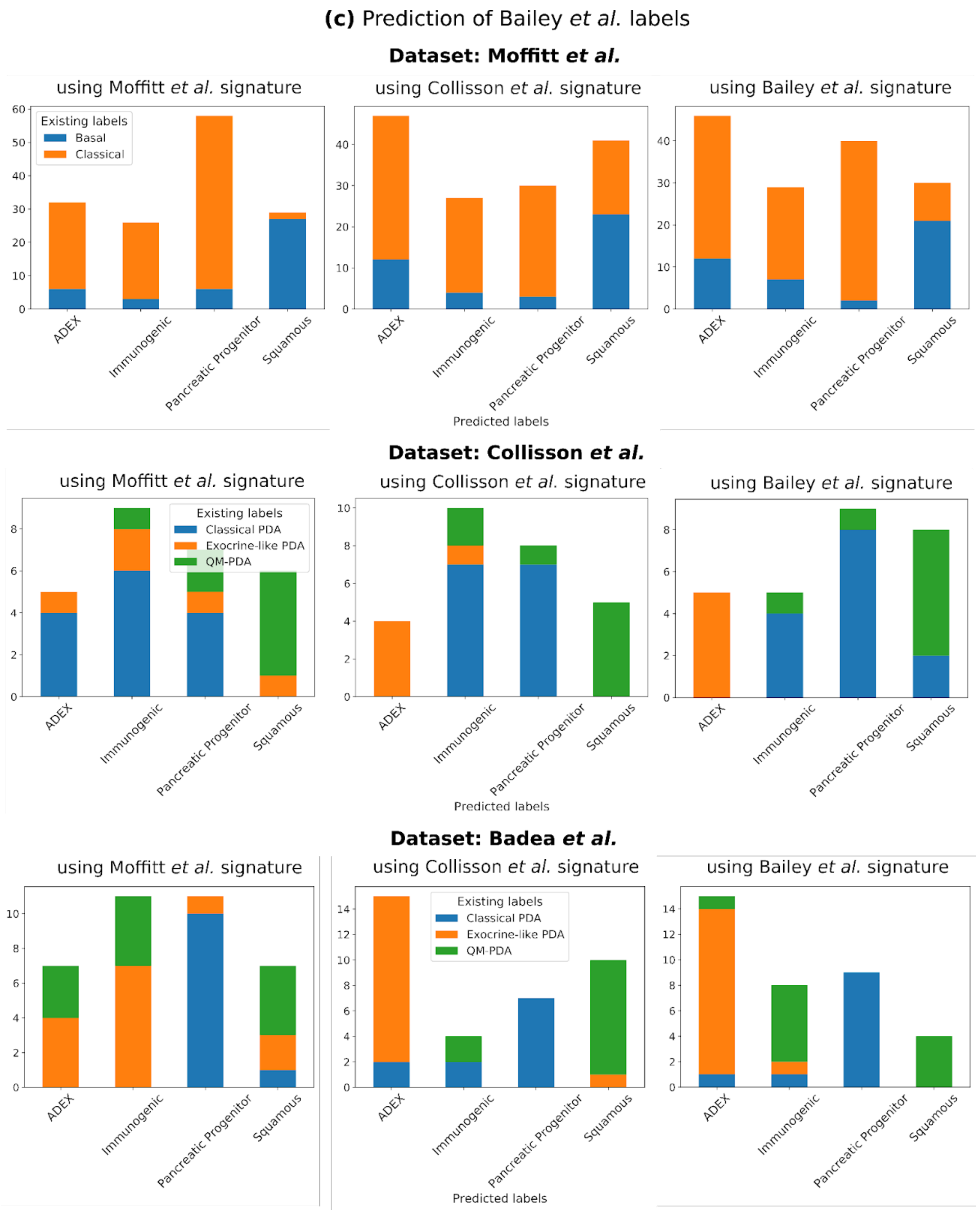


**Figure S3.** Comparison between predicted subtypes and existing subtypes assigned by the studies. For each classification model and for each dataset whose subtypes are known is shown the composition of the predicted subtypes, on the x-axis, with respect to the real ones. (a) Models trained on Moffitt *et al.* dataset are used to classify samples into Basal and Classical subtype using signature from Moffitt *et al.* (left figure), secondly

Collisson *et al.* (central figure) and lastly Bailey *et al.* (right figure). Validation dataset considered for this comparison are the ones with existing subtype labels, excluding the same dataset used for validation. Considering one dataset and a time, we can check whether the subtype assigned to each sample through classification corresponds to the existing subtype by looking at the composition of the stacked bars in the figure. (b) and (c) are the same as (a) but showing Collisson et al. and Bailey et al. predicted subtypes, respectively.

#### Figure S4

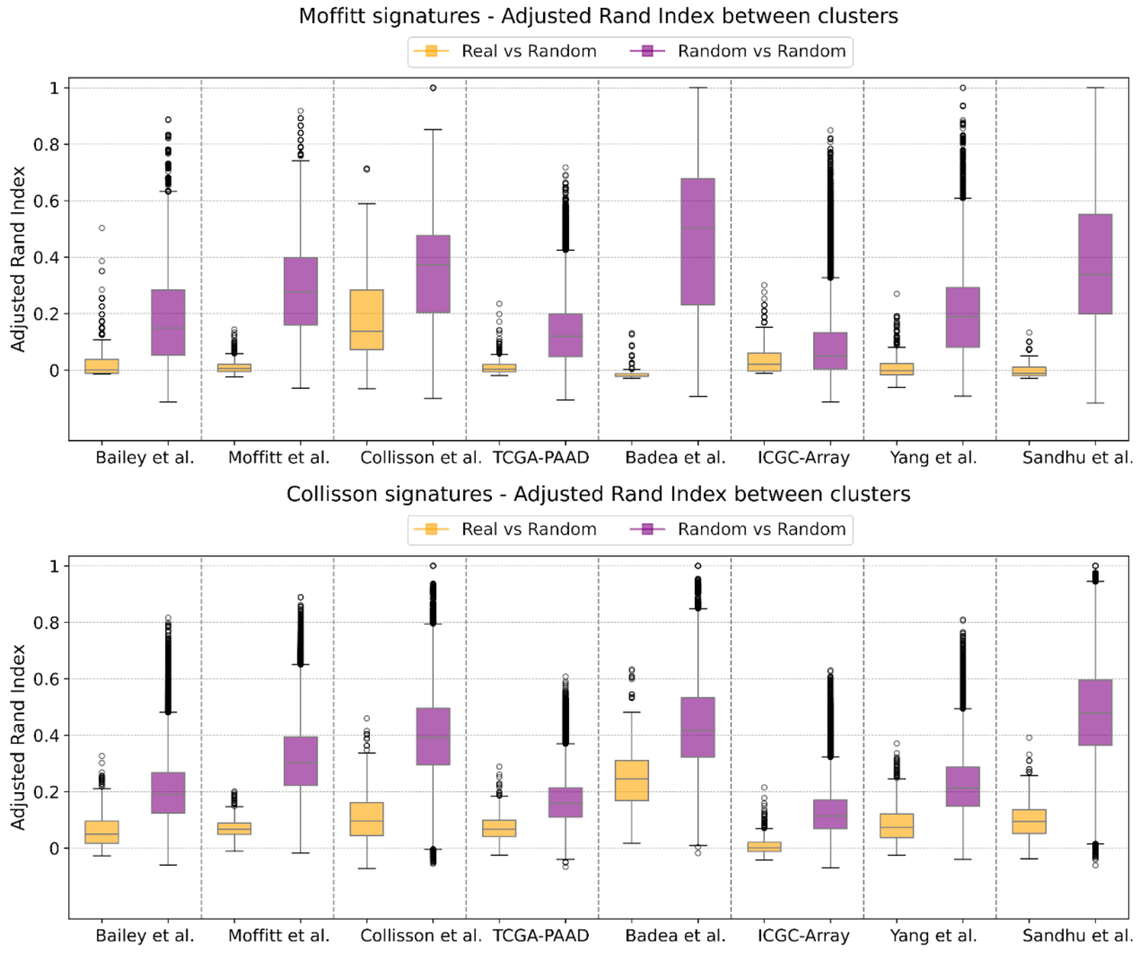

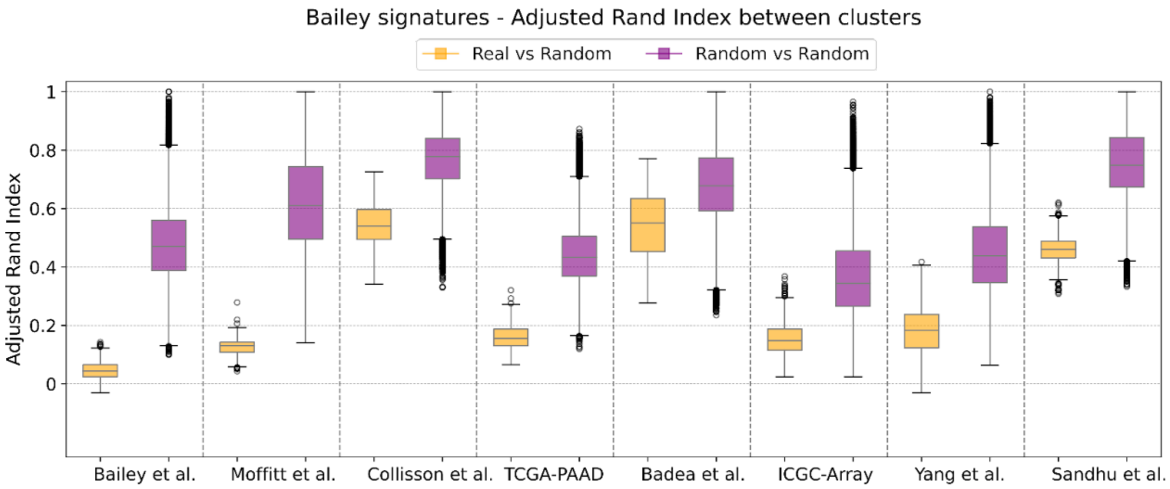


**Figure S4.** Adjusted Rand Index (ARI) is used for evaluating the clustering robustness of the signatures. Clusters derived using Moffitt *et al.* (top), Collisson *et al.* (center) and Bailey *et al.* (bottom) signatures are compared with the ones identified when employing random genes of the same size of the signature (Real vs Random). ARI is also used for a pair-wise comparison between clusters deriving from the use of random gene sets (Random vs Random).

### Figure S5

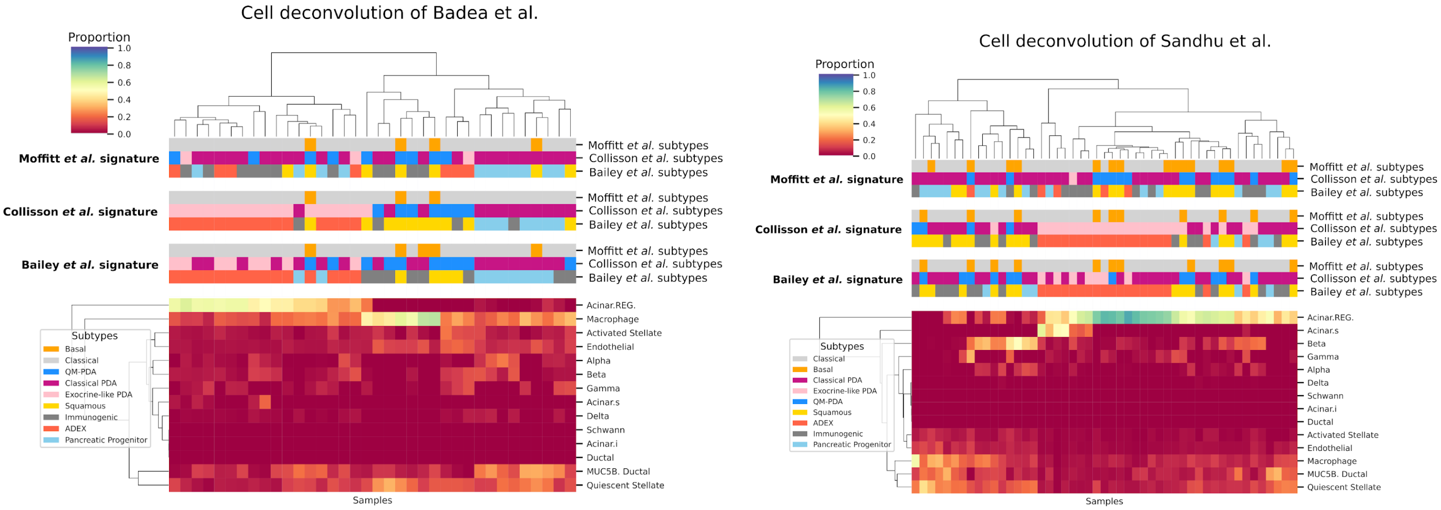

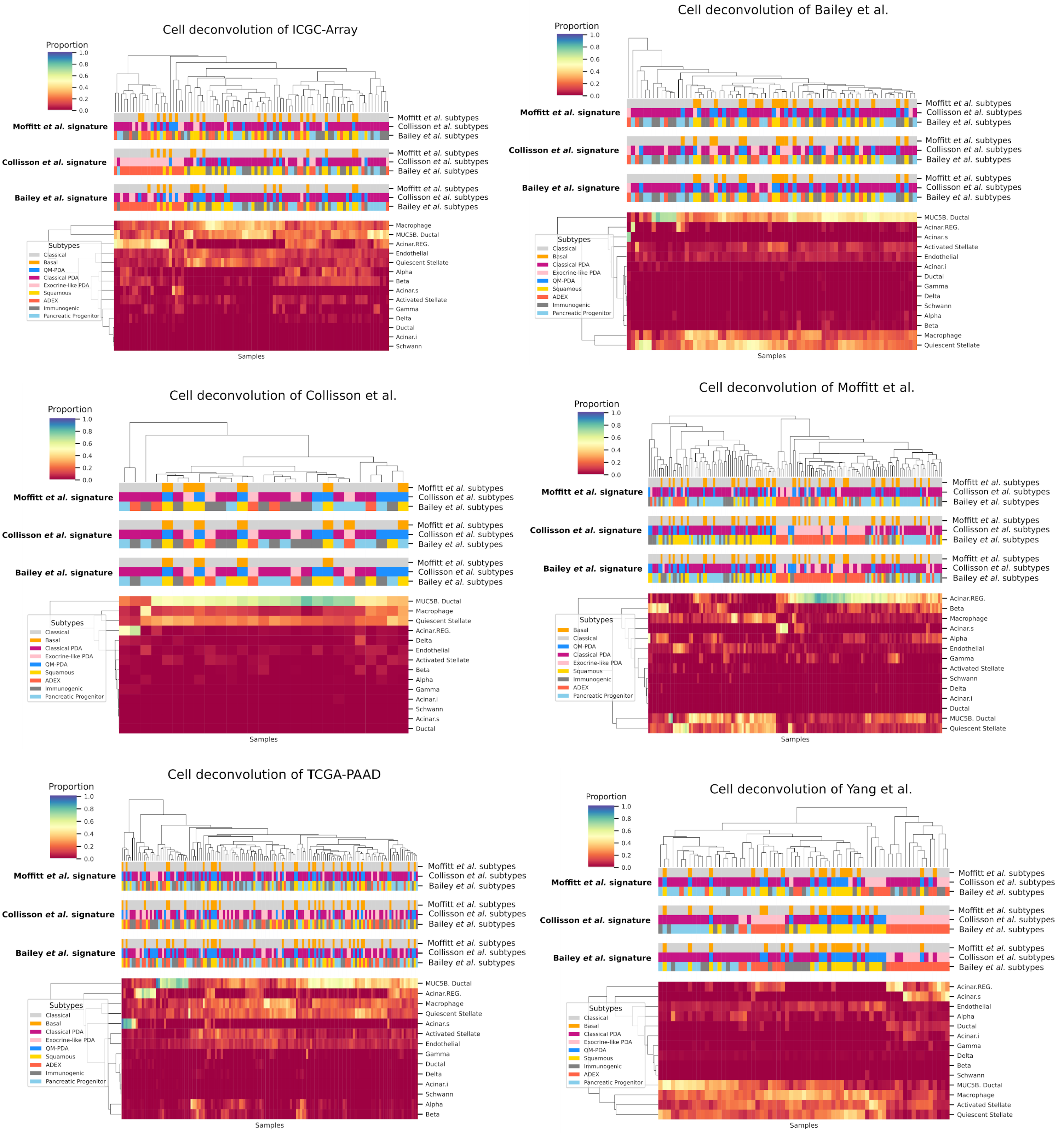


**Figure S5.** Hierarchical clustering of the inferred proportion of pancreatic cell types enriched in the eight cohorts. For each dataset are shown the classes predicted by the nine predictor models.

Figure S6

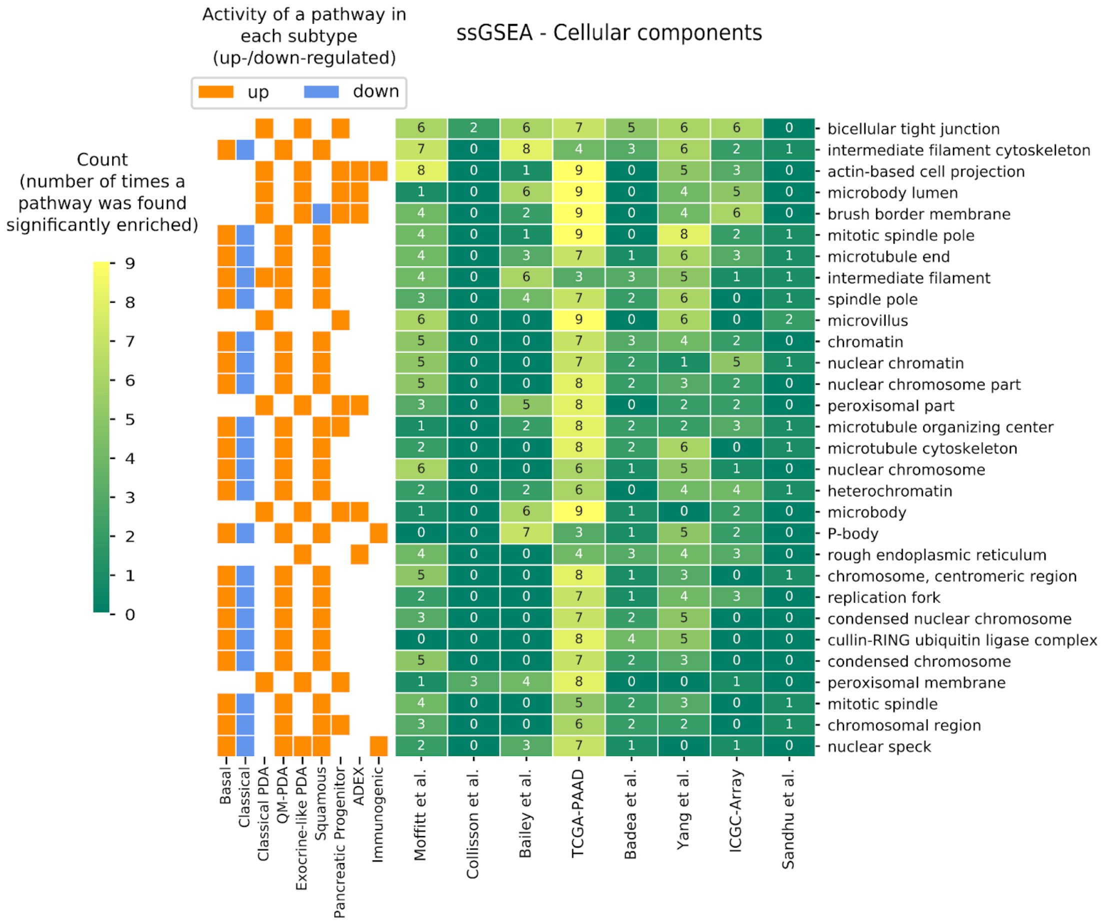

Activity of a pathway in each subtype (up-/down-regulated)
up
down
ssGSEA - Cellular components
Count (number of times a pathway was found significantly enriched)
9
8
7
6
5
4
3
2
1
0
bicellular tight junction
intermediate filament cytoskeleton
actin-based cell projection
microbody lumen
brush border membrane
mitotic spindle pole
microtubule end
intermediate filament
spindle pole
microvillus
chromatin
nuclear chromatin
nuclear chromosome part
peroxisomal part
microtubule organizing center
microtubule cytoskeleton
nuclear chromosome
heterochromatin
microbody
P-body
rough endoplasmic reticulum
chromosome, centromeric region
replication fork
condensed nuclear chromosome
cullin-RING ubiquitin ligase complex
condensed chromosome
peroxisomal membrane
mitotic spindle
chromosomal region
nuclear speck
Basal
Classical
Classical PDA
QM-PDA
Exocrine-like PDA
Squamous
Pancreatic Progenitor
ADEX
Immunogenic
Moffitt et al.
Collisson et al.
Bailey et al.
TCGA-PAAD
Badea et al.
Yang et al.
ICGC-Array
Sandhu et al.

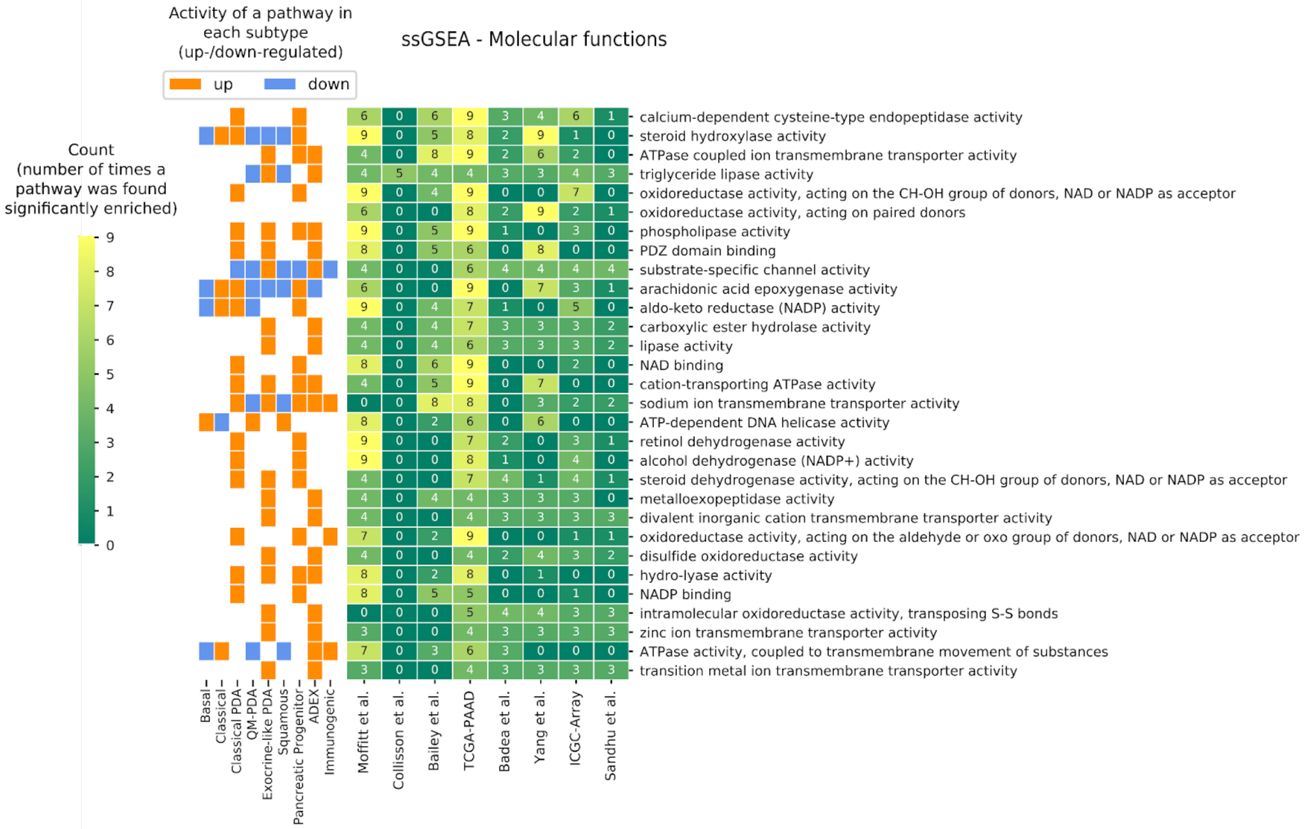
Activity of a pathway in each subtype (up-/down-regulated)
up
down
ssGSEA - Molecular functions
Count (number of times a pathway was found significantly enriched)
calcium-dependent cysteine-type endopeptidase activity
steroid hydroxylase activity
ATPase coupled ion transmembrane transporter activity
triglyceride lipase activity
oxidoreductase activity, acting on the CH-OH group of donors, NAD or NADP as acceptor
oxidoreductase activity, acting on paired donors
phospholipase activity
PDZ domain binding
substrate-specific channel activity
arachidonic acid epoxygenase activity
aldo-keto reductase (NADP) activity
carboxylic ester hydrolase activity
lipase activity
NAD binding
cation-transporting ATPase activity
sodium ion transmembrane transporter activity
ATP-dependent DNA helicase activity
retinol dehydrogenase activity
alcohol dehydrogenase (NADP+) activity
steroid dehydrogenase activity, acting on the CH-OH group of donors, NAD or NADP as acceptor
metalloexopeptidase activity
divalent inorganic cation transmembrane transporter activity
oxidoreductase activity, acting on the aldehyde or oxo group of donors, NAD or NADP as acceptor
disulfide oxidoreductase activity
hydro-lyase activity
NADP binding
intramolecular oxidoreductase activity, transposing S-S bonds
zinc ion transmembrane transporter activity
ATPase activity, coupled to transmembrane movement of substances
transition metal ion transmembrane transporter activity
Basal
Classical
Classical PDA
QM-PDA
Exocrine-like PDA
Squamous
Pancreatic Progenitor
ADEX
Immunogenic
Moffitt et al.
Collisson et al.
Bailey et al.
TCGA-PAAD
Badea et al.
Yang et al.
ICGC-Array
Sandhu et al.

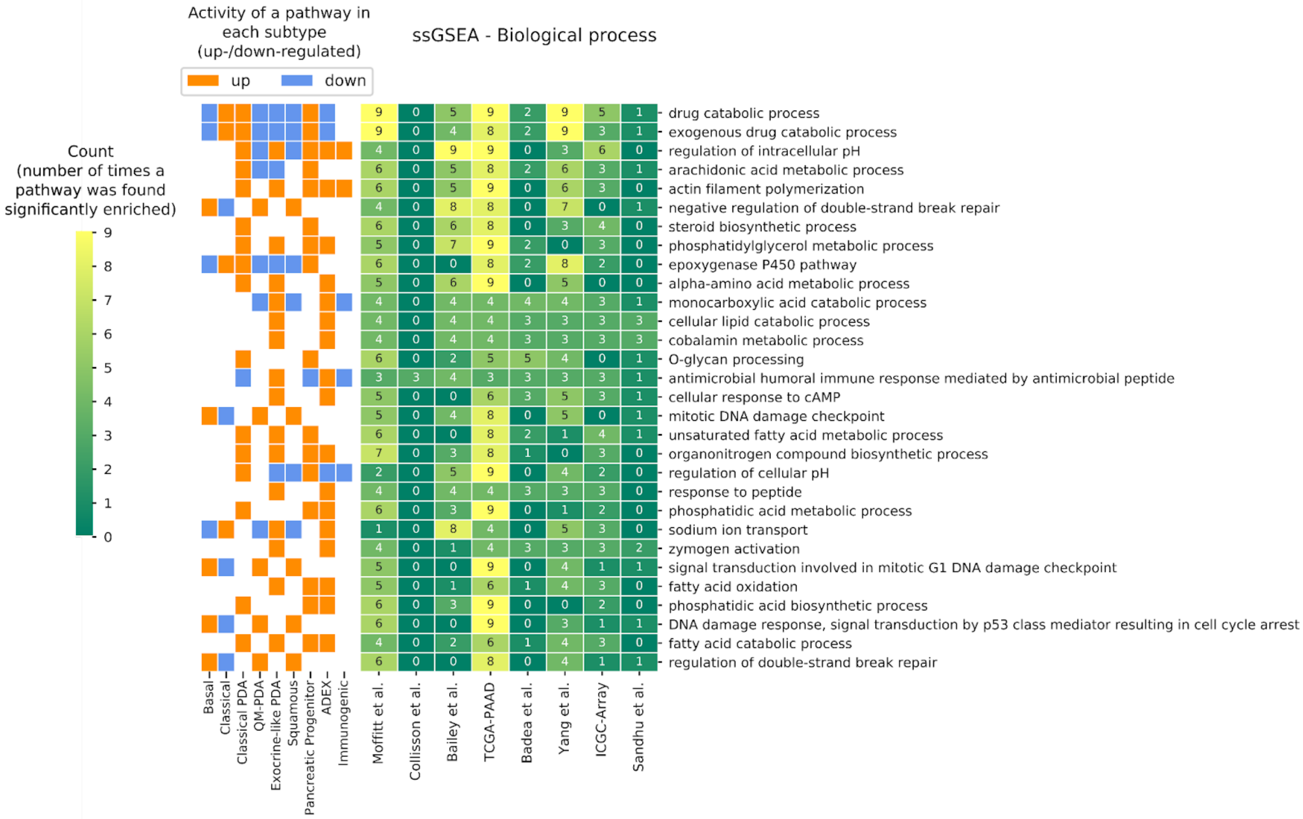
Activity of a pathway in each subtype (up-/down-regulated)
up
down
ssGSEA - Biological process
Count (number of times a pathway was found significantly enriched)
9
8
7
6
5
4
3
2
1
0
drug catabolic process
exogenous drug catabolic process
regulation of intracellular pH
arachidonic acid metabolic process
actin filament polymerization
negative regulation of double-strand break repair
steroid biosynthetic process
phosphatidylglycerol metabolic process
epoxygenase P450 pathway
alpha-amino acid metabolic process
monocarboxylic acid catabolic process
cellular lipid catabolic process
cobalamin metabolic process
O-glycan processing
antimicrobial humoral immune response mediated by antimicrobial peptide
cellular response to cAMP
mitotic DNA damage checkpoint
unsaturated fatty acid metabolic process
organonitrogen compound biosynthetic process
regulation of cellular pH
response to peptide
phosphatidic acid metabolic process
sodium ion transport
zymogen activation
signal transduction involved in mitotic G1 DNA damage checkpoint
fatty acid oxidation
phosphatidic acid biosynthetic process
DNA damage response, signal transduction by p53 class mediator resulting in cell cycle arrest
fatty acid catabolic process
regulation of double-strand break repair
Basal
Classical
Classical PDA
QM-PDA
Exocrine-like PDA
Squamous
Pancreatic Progenitor
ADEX
Immunogenic
Moffitt et al.
Collisson et al.
Bailey et al.
TCGA-PAAD
Badea et al.
Yang et al.
ICGC-Array
Sandhu et al.

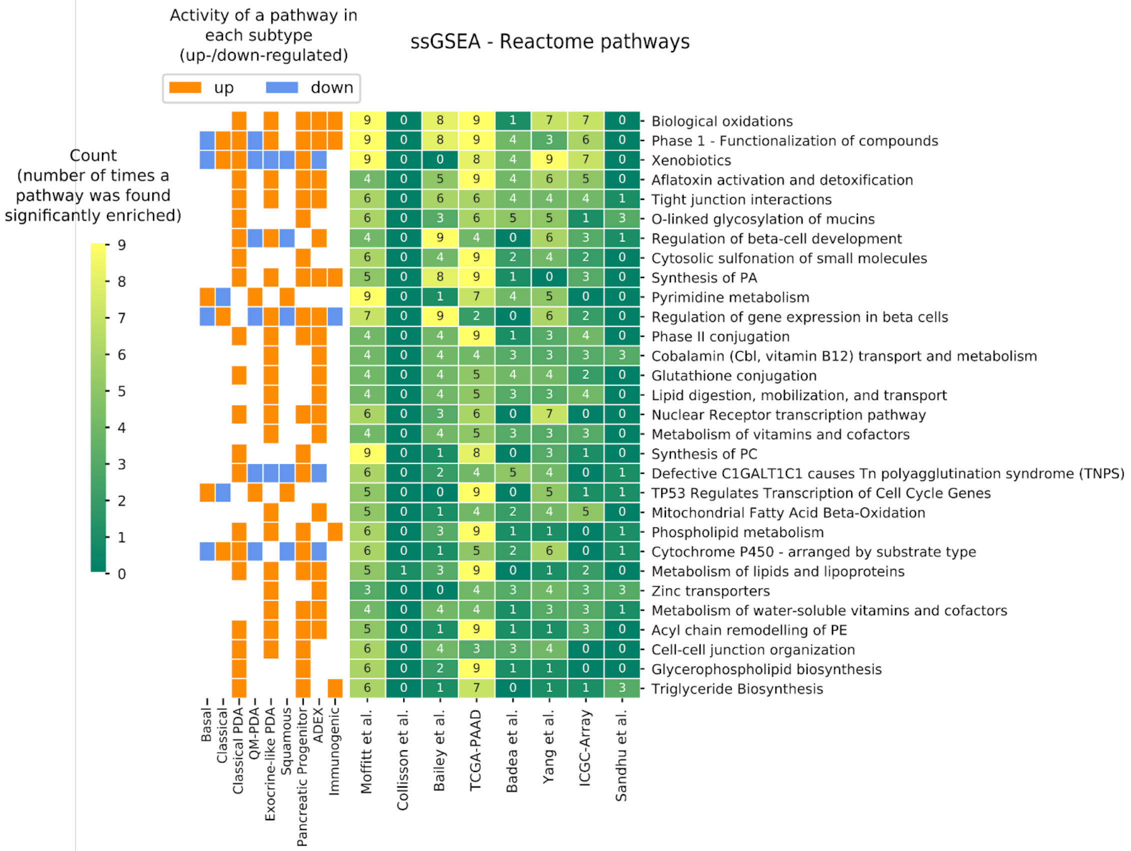


**Figure S6.** ssGSEA of the eight cohorts. Enrichment scores of terms are compared between predicted subtypes and across cohorts and the top 30 most significant are counted and shown. For each term we know whether it was found up or down regulated and in which subtype.
